# The three YTHDF paralogs and VIRMA are the major tumor drivers among the m^6^A core genes in a pan-cancer analysis

**DOI:** 10.1101/2024.06.13.598899

**Authors:** Eliana Destefanis, Denise Sighel, Davide Dalfovo, Riccardo Gilmozzi, Francesca Broso, Andrea Cappannini, Janusz M. Bujnicki, Alessandro Romanel, Erik Dassi, Alessandro Quattrone

## Abstract

N^6^-methyladenosine (m^6^A) is the most abundant internal modification in mRNAs. Despite accumulating evidence for the profound impact of m^6^A on cancer biology, there are conflicting reports that alterations in genes encoding the m^6^A machinery proteins can either promote or suppress cancer, even in the same tumor type. Using data from The Cancer Genome Atlas, we performed a pan-cancer investigation of 15 m^6^A core factors in nearly 10,000 samples from 31 tumor types to reveal underlying cross-tumor patterns. Altered expression, largely driven by copy number variations at the chromosome arm level, results in the most common mode of dysregulation of these factors. YTHDF1, YTHDF2, YTHDF3, and VIRMA are the most frequently altered factors and the only ones to be uniquely altered when tumors are grouped according to the expression pattern of the m^6^A factors. These genes are also the only ones with coherent, pan-cancer predictive power for progression-free survival. On the contrary, METTL3, the most intensively studied m^6^A factor as a cancer target, shows much lower levels of alteration and no predictive power for patient survival. Therefore, we propose the non-enzymatic YTHDF and VIRMA genes as preferred subjects to dissect the role of m^6^A in cancer and as priority cancer targets.

**Graphical abstract:** 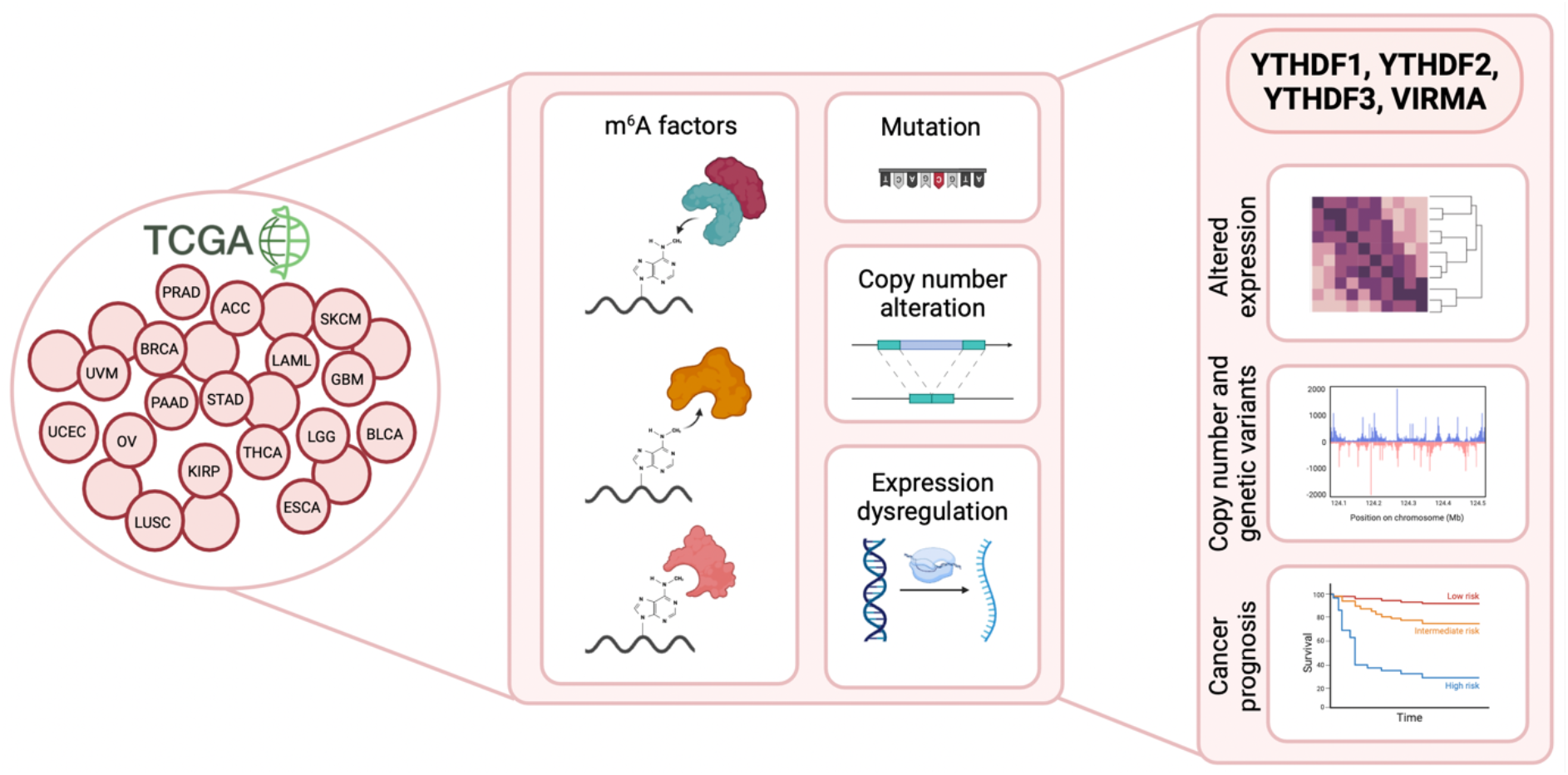

## Introduction

N^6^-methyladenosine (m^6^A) is the most prevalent internal modification in mRNAs and long non-coding RNAs (lncRNAs) (1). It is enriched at stop codons and in 3’ untranslated regions (3’ UTR), with a well conserved DRACH (D = A, G, U; R = A, G; H = A, C, U) consensus motif across eukaryotes (1, 2).

m^6^A is a dynamic and reversible modification regulated by writer, eraser and reader proteins (3). Specifically, the m^6^A writer proteins, which introduce the modification, include the m^6^A-METTL catalytic complex (MAC: METTL3 and METTL14) and the m^6^A-METTL associated complex (MACOM: WTAP, VIRMA - also known as KIAA1429, RBM15 or RBM15B, ZC3H13, HAKAI - also known as CBLL1) (4). The FTO and ALKBH5 proteins are the demethylases, responsible for removing the m^6^A modification (5, 6). Finally, the reader proteins include the cytoplasmic YTHDF1-3 and the nuclear YTHDC1-2 proteins, which recognize N^6^-methylated adenine through the YTH domain (1, 7–12). In addition, other non-YTH RNA-binding proteins such as eIF3a, HNRNPC/G, HNRNPA2B1, IGF2BP1/2/3, and FMRP have been shown to bind m^6^A in a sequence-dependent manner, although their roles are not exclusively related to m^6^A (13–19). By detecting changes in m^6^A modification status, reader proteins can influence downstream cellular processes such as transcription, splicing, localization, translation, and degradation (20).

Associations between dysregulated m^6^A levels and various complex disorders, including cancer, have been extensively studied and experimentally validated (21). In cancer, m^6^A dynamics have been implicated in tumor initiation, maintenance, and metastasis (22). Indeed, aberrant m^6^A levels, including both increased and decreased levels, have been shown to promote tumorigenesis by affecting different cellular processes (23, 24). Consistent with these findings, alterations in the same m^6^A machinery proteins have been shown to either promote or suppress cancer initiation and/or progression, even within the same tumor type (25). For example, reduced METTL3 or METTL14 expression has been demonstrated to promote glioblastoma tumorigenesis, while METTL3 overexpression or FTO inhibition has been shown to suppress glioblastoma stem cell growth and self-renewal in vitro and in vivo (26).

Conversely, another study has revealed that METTL3 expression and high m^6^A levels are crucial for maintaining the glioblastoma stem cells pool (27). Similarly, in non-small cell lung cancer (NSCLC), ALKBH5 has been demonstrated to both promote (28, 29) and inhibit (30) tumor progression by targeting different downstream genes. In addition, m^6^A factors with opposite enzymatic roles have also shown oncogenic effects within the same tumor type. For instance, in acute myeloid leukemia (AML), METTL3 has been found to be abundantly expressed and to maintain the undifferentiated leukemic phenotype by increasing m^6^A modifications in key AML oncogenes, and its depletion or pharmacological inhibition leads to proliferation reduction and differentiation promotion (31–33). However, conflicting results have been reported, as FTO has also been shown to be highly expressed in AML, particularly in mixed-lineage leukemia-rearranged samples, where it promotes leukemogenesis by reducing m^6^A levels on transcripts such as ASB2 and RARA (34) and its inhibition can be exploited therapeutically (35–37). A similar scenario has been observed in other tumors, such as bladder, lung, and pancreatic cancers (38–44)

Given this complex and controversial landscape, a comprehensive analysis of alterations in the genes encoding for the m^6^A machinery across various cancer types could provide an unbiased picture of their potential roles as oncogenes or tumor suppressors. Here, we performed a computational investigation of alterations in the m^6^A machinery genes across 31 tumor studies from the Tumor Cancer Genome Atlas (TCGA). We found that altered expression due to arm-level chromosomal imbalance is the most predominant type of dysregulation and showed cross-tumor profiles of m^6^A factor expression. Unexpectedly, YTHDF1, YTHDF2, YTHDF3, and VIRMA, rather than the intensively studied enzymatic m^6^A proteins, emerged as the most frequently dysregulated factors, and their alterations significantly associated with patient prognosis.

## Materials and Methods

### m^6^A factors

Among the genes encoding the m^6^A machinery, 15 were selected. In particular, METTL3, METTL14, RBM15, RBM15B, WTAP, VIRMA, ZC3H13, and HAKAI were considered to be part of the multi-subunit writer complex (4). Furthermore, among the reader proteins, YTHDF1, YTHDF2, YTHDF3, YTHDC1, and YTHDC2 were chosen as being direct interactors with the m^6^A methylation. Finally, the erasers ALKBH5 and FTO were also included (3).

### Tumor sample datasets

Alteration data of tumor samples were downloaded from the Tumor Cancer Genome Atlas (TCGA). TCGA samples for which copy number alteration, mutation and gene expression data were all available, or for which only expression and copy number alteration data could be retrieved, were included, resulting in 6956 and 9327 samples, respectively, across 31 cancer types (**Supplementary Table S1**). Data for 15 m^6^A factors were obtained for these samples using the cBioPortal R package, *cgdsr* (v1.3.0) (45, 46). GISTIC 2.0 (47) profiles were selected for copy number alterations, rna_seq_v2_mrna_median_Zscores profiles were chosen for gene expression, and all the mutations were considered. In addition, data were retrieved for the genes annotated as oncogenes and/or tumor suppressors in OncoKB (541 cancer drivers, https://www.oncokb.org/). For TCGA samples, tumor stages, grades, and molecular subtypes were retrieved from the *cgdsr* and *TCGAbiolinks* R packages (v2.12.6) (48). Tumor samples mRNA expression levels (TPM transcript per kilobase million) were obtained from Xena RSEM Toil Recomputed data (https://xena.ucsc.edu/) (49).

### Computation of the sample alteration scores

Copy number alteration (CNA), mutation, and gene expression data were retrieved for all the investigated genes in all tumor samples. Low-level CNAs were collected for a gene when the absolute relative GISTIC score was 1, while a high-level CNA was recorded when the absolute relative GISTIC score was 2. Similarly, a differential expression was reported for those genes with an absolute z-score >= 2. Gene mutations were counted when they occurred. The final alteration score was defined as 1 for overexpression, amplification, or mutation, -1 for downregulation or deep deletion, and 0 if the gene remained unaltered in that sample.

To assess the significance of the alteration frequencies for the 15 m^6^A factors, an empirical p-value was computed by first comparing the total alteration frequencies of the m^6^A factors with those of 15 randomly selected genes (out of 18421 total genes) and then with those of 15 randomly selected cancer driver genes (out of 541 total genes). In both cases, a 1000 iteration bootstrap was applied. Similarly, an empirical p-value was calculated by comparing the upregulation frequencies of the 15 m^6^A factors with the upregulation frequencies of 15 randomly selected genes and cancer driver genes with a 1000-iteration bootstrap.

For the expression and copy number alterations, paired Wilcoxon tests were employed to compare the frequencies of upregulation/amplification and downregulation/deep deletion of the m^6^A factors across all tumor samples. Additionally, paired Wilcoxon tests were performed for each m^6^A factor individually, comparing their upregulation/amplification and downregulation/deep deletion frequencies within each tumor type and vice versa for each tumor type with respect to the upregulation/amplification and downregulation/deep deletion frequencies of each m^6^A factor.

### Analysis of m^6^A factor mutations

Tumor sample mutation data were obtained by downloading publicly available data from the TCGA data analysis center (GDAC) repository (https://gdac.broadinstitute.org/). Mutations were annotated using the Variant Effect Predictor (VEP) tool using the GRCh37.p13 assembly.

### Identification of m^6^A subtypes

Tumor samples altered in at least one m^6^A factor (6092) were clustered using a non-negative matrix factorization (NMF) algorithm. A matrix containing the binarized version of the alteration score (1 if an expression alteration occurs for a factor in a sample, 0 otherwise) for each sample was used for the clustering using the *NMF* R package (v0.23.0) (50). The number of factorization ranks was estimated and then fourteen was selected. The algorithm was run 100 times, and the Brunet method was applied (51).

To identify representative m^6^A machinery genes for each rank, the percentage of samples altered in each m^6^A factor was computed for each rank, and then z-score standardized. A threshold of 1 was applied to the z-scores to select the representative genes.

The comparison between the expression of m^6^A factors (TPM, Xena portal) in the selected and other classes was performed using the Wilcoxon test with a 0.05 p-value cutoff.

### Enrichment analysis

Enrichment analyses were performed using Fisher exact tests. A Benjamini-Hochberg correction was applied to the p-values, and a 0.05 cutoff was set. Only terms with odds ratios >= 3 were taken into consideration. Pathway enrichment was performed with *clusterProfiler* (v3.14.0) and *ReactomePA* (v1.30.0), using KEGG and Reactome annotations (52, 53).

### mRNA-protein analysis

Protein levels and corresponding mRNA levels were obtained from a compendium dataset of 2002 primary tumors from 14 cancer types and 17 studies (54). For each tissue and m^6^A factor, the protein expression levels were compared between the samples with upregulated/downregulated mRNA (absolute z-score >= 2) and the remaining samples.

### eQTL analysis

Genotype calls generated from Affymetrix SNP Array 6.0 used in the heritability analysis were retrieved from the TCGA legacy archive (portal.gdc.cancer.gov/legacy-archive). Stringent quality control measures were applied to the SNP genotyping data. Only autosomal SNP were considered. SNPs and individuals with call rates <90% were excluded. Multi-allelic SNPs and SNPs with minor allele frequencies <1% or Hardy-Weinberg equilibrium test p-values < 10^-6^ were discarded. Overall, genotype calls of 842,108 SNPs across 10,755 TCGA samples were considered.

Estimates of genome-wide heritability of the cancer-associated m^6^A factors were performed using GCTA (v1.94.1), which estimates the proportion of variance in a phenotype explained by all SNPs (55). The GCTA GREML approach was used with the default average information (AI) algorithm to run REML iterations. The genetic relatedness matrix (GRM) was calculated as a measure of the genetic similarity for unrelated individuals (GRM < 0.05) and then compared to the similarity of the selected m^6^A factors to compute the contribution of the genotypic variance to overall phenotypic variance, V(Genotype)/V(Phenotype).

Cis-eQTLs, survival-associated eQTLs, and GWAS-associated eQTLs for all the TCGA tumors were retrieved from the PancanQTL database (http://bioinfo.life.hust.edu.cn/PancanQTL/) (56). eQTLs annotated to the selected m^6^A factors were retrieved, and the respective positive or negative beta values (effect size of SNP on gene expression) were considered.

Linkage disequilibrium (LD) analyses were performed for each selected m^6^A factor. LD was computed using 1000 Genomes Project individual genotypes using *ldsep* R package (v2.1.5), which implements a maximum likelihood approach to estimate LD measures (57). Specifically, LD was calculated between each pair of associated SNPs of the annotated eQTLs. All SNPs in strong LD were aggregated in groups. Strong LD was defined as R^2^ > 0.8. The number of strong LD groups for each tumor type was represented using *circlize* R package (v0.4.15) (58).

### Nearest cancer driver genes identification

For each m^6^A factor, the nearest cancer driver gene was extracted (out of 541 cancer drivers downloaded from OncoKB, https://www.oncokb.org/), considering genes laying on the same chromosomal arm and with a maximum 1 Mb of distance. GISTIC profiles were retrieved for each gene in all the tumor samples. Gain and amplification events were recorded for GISTIC scores equal to 1 and 2, respectively. Oppositely, deletion and deep deletion were represented by -1 and -2 scores, respectively. The concordant presence of copy number alterations on the m^6^A factor and its near cancer driver gene was explored considering samples belonging to the enriched tumor types and with expression alterations in the respective m^6^A factor.

### m^6^A factors and cancer drivers co-occurrent and mutually exclusive interactions

identification of co-occurrent and mutually exclusive alterations was performed using the SELECT algorithm (v1.6) (59). A binary matrix with the m^6^A factors and cancer driver genes positive and negative expression or copy number alteration scores of tumor samples with at least one altered gene was used as input. The p-values of the weighted version of the mutual information (wMI) were converted to negative values for the mutually exclusive gene pairs and transformed to 1 when comparing the same gene. Interactions that met both the False Discovery Rate (FDR) cutoff and the Average Sum Correction (ASC) effect size threshold were considered.

### Survival analysis

TCGA survival outcome data were obtained from the standardized datasets published by Liu et al. (60). The progression-free interval (PFI) endpoint was used in the pan-cancer analysis and samples without PFI survival time reported were filtered out. For single tumor survival analysis, the overall survival (OS) was used when more endpoint possibilities were available; otherwise, suggested endpoints between PFI and disease-specific survival (DSS) were applied. The PCPG samples were removed as no suitable endpoints were available, as suggested by Liu et al. (60).

Survival analyses were performed on different settings. In one setting, samples with expression alterations in one m^6^A factor were compared to those without. For this analysis, all 9327 tumor samples were considered. In another experimental setting, survival analyses were performed by considering only the classes presenting an altered m^6^A factor. For this analysis, samples altered in at least one m^6^A factor (6092 samples) were included, and samples of a class were compared to samples of the other classes.

The analyses were performed in a pan-cancer mode, considering all the tumor types, or in single-tumor mode, considering each tumor type separately. The positive and negative alterations scores were considered separately.

Cox regression-hazard model was applied in all survival analyses. Survival analyses in the pan-cancer mode were performed with a multivariate cox regression analysis using the tumor type, age and sex as covariates. Instead, the survival analyses of the single tumor-mode were performed with a multivariate cox regression analysis using the age and sex as covariates. All the analyses were performed using the *survival* and *survimer* R packages (v2.44-1.1, v0.4.6) (61, 62).

Those comparisons where at least 10 samples were present, in both altered and control subsets of patients, were taken into consideration. In the pan-cancer analysis a 0.1 p-value cutoff was applied. In the single tumor analysis, Benjamini-Hochberg correction was computed for each tumor type across the tested conditions (m^6^A factors/classes) and a 0.1 adjusted p-value cutoff was applied.

A quantile normalization was applied on the PFI endpoints of samples of each tumor type, separately. Survival analyses performed on the samples with expression alterations in one m^6^A factor compared to those without, were run also with the normalized PFI times. The results obtained with the standard and normalized PFI times were transformed in 1/-1 and 0 if significant or not significant, respectively, with the sign corresponding to the hazard ratio. The consistency between the two methods was calculated by computing the percentage of concordant and opposite results on the total number of tests.

## Results

### 1. Dysregulated expression of the YTHDF and VIRMA genes is the most frequent cross-tumor alteration among the m^6^A core factors

To investigate the pan-cancer profile of the factors comprising the m^6^A machinery, we selected the core 15 m^6^A writer, eraser and reader genes (writers: METTL3, METTL14, VIRMA, WTAP, ZC3H13, HAKAI and RBM15/15B; readers: YTHDC1/2, YTHDF1/2/3; erasers: ALKBH5 and FTO) and analyzed their variation in terms of mutations, differential expressions, and copy number alterations (CNAs) in 6956 TGCA cancer samples belonging to 31 tumor types (**Figure 1A**, **Supplementary Table S1**). We assumed differential expression significant only for absolute z-scores greater than 2, and retained CNAs only at high gene copy levels, which are supposed to represent amplifications and homozygous deletions (45, 46). Indeed, the pattern of CNAs at low gene copy levels, presumed to represent gains and heterozygous deletions, was rather widespread (**Figure S1A**) (63). The majority of samples (70.6%) scored to be altered for either mutation, expression, or CNA in at least one of the selected m^6^A core factors, with altered expression (63.9%) being predominant (**Figure 1B**). Altered samples belong to all tumor types considered, although to different extents (**Figure S1B, Supplementary Tables S2-4**), and the majority are altered for only one of the 15 m^6^A core factors (42.8%) (**Figure S1C**). While writers and readers are more altered than erasers, accounting for about three times as many samples with at least one altered gene (**Figure 1C**), the average normalized frequency of alteration is the same (10%, 12% and 9%, respectively). Looking at the individual factors, the percentage of alterations ranges from 21.4% for VIRMA to 6.1% for RBM15 (**Figure 1C**, **Supplementary Tables S2-4**).

**Figure 1.**
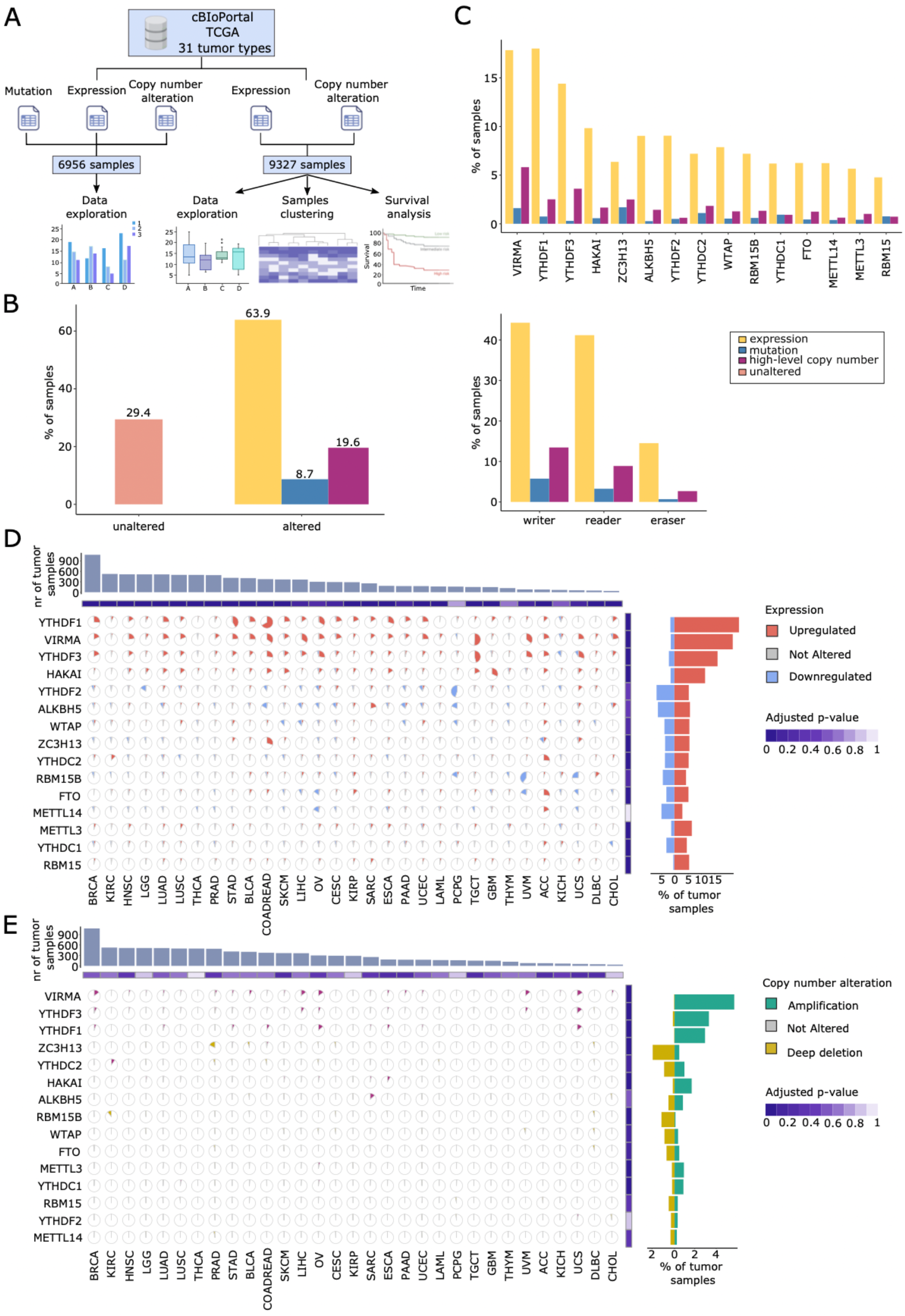
Expression dysregulation is the most frequent alteration of m^6^A factors across 31 tumor types. A) Graphical representation of the types of alteration considered from the TCGA dataset, the corresponding number of samples, and downstream analyses. B) The plot shows the percentage of samples altered in m^6^A factors stratified by presence/absence of high-level copy number, mutation, and/or expression alterations. C) The plot shows the percentage of samples altered in m^6^A factors stratified by m^6^A factor roles (up) and factors (bottom). D) Pie charts display the percentage of tumor samples exhibiting changes in expression levels (upregulation: z-score => 2; downregulation: z-score <= -2) categorized by m^6^A factors and tumor types. The number of samples of each tumor type is depicted at the top of the plot. The percentage of samples displaying upregulation and downregulation for each m^6^A factor is illustrated on the right side of the chart. Paired Wilcoxon test was performed to compare the frequencies of upregulation and downregulation in each tumor type and for each m^6^A factor (adjusted p-value <= 0.05). E) Pie charts display the percentage of tumor samples exhibiting changes in copy number events (amplification: GISTIC score = 2; deep deletion: GISTIC score = -2) categorized by m^6^A factors and tumor types. The number of samples of each tumor type is depicted at the top of the plot. The percentage of samples displaying amplification and deep deletion for each m^6^A factor is illustrated on the right side of the chart. A paired Wilcoxon test was performed to compare the frequencies of amplification and deep deletion in each tumor type and for each m^6^A factor (adjusted p-value <= 0.05). Cancer type abbreviations: ACC, Adrenocortical Cancer; BLCA, Bladder Cancer; BRCA, Breast Cancer; CESC, Cervical Cancer; CHOL, Cholangiocarcinoma; COADREAD, Colorectal Cancer; DLBC, Large B-cell Lymphoma; ESCA, Esophageal Cancer; GBM, Glioblastoma; HNSC, Head and Neck Cancer; KICH, Kidney Chromophobe; KIRC, Kidney Clear Cell Carcinoma; KIRP, Kidney Papillary Cell Carcinoma; LAML, Acute Myeloid Leukemia; LGG, Lower Grade Glioma; LIHC, Liver Cancer; LUAD, Lung Adenocarcinoma; LUSC, Lung Squamous Cell Carcinoma; OV, Ovarian Cancer; PAAD, Pancreatic Cancer; PCPG, Pheochromocytoma & Paraganglioma; PRAD, Prostate Cancer; SARC, Sarcoma; SKCM, Melanoma; STAD, Stomach Cancer; TGCT, Testicular Cancer; THCA, Thyroid Cancer; THYM, Thymoma; UCEC, Uterine Corpus Endometrial Carcinoma; UCS, Uterine Carcinosarcoma; UVM, Ocular melanoma.

Remarkably, the m^6^A core factors have an overall higher pan-cancer alteration frequency not only when compared to 15 randomly selected genes (p-value < 10^-3^), but also when compared to 15 randomly selected cancer driver genes (p-value = 0.03; n=541 cancer driver genes from OncoKB) with a 1000 iteration bootstrap.

We next focused on the individual occurrences of mutations, differential expression and CNAs, to gain insight into their functional involvement in tumorigenesis. The total number of point mutations in the 15 m^6^A core factor genes was only 758, predominantly missense (76%), with an average mutation frequency of 0.7%, lower than in the 541 cancer driver genes (1.1%) (**Figure S1D**). Moreover, by investigating the mutation landscape of the 15 m^6^A factors using available classifications of cancer driver genes, we found that none were classified as cancer drivers or weak drivers (64–67), apart from RBM15, which emerged as a driver specifically in head and neck squamous cell carcinoma (HNSC) and lung adenocarcinoma (LUAD) (66). RBM15 has been reported as a potential cancer gene also for its involvement in the RBM15-MKL1 fusion in AML (68, 69). Analysis of the differential expression of the 15 m^6^A core factors (9327 TCGA samples, **Figure 1A**) revealed that most factors (11/15) are more frequently upregulated than downregulated (paired Wilcoxon test p-value = 0.003) (**Figure 1D**). Notably, the core factors show a higher mean upregulation when compared to 15 randomly selected genes and to 15 randomly selected oncogenes (1000 iterations bootstrap, p-value < 10^-3^ and p-value=0.005, respectively; n=286 oncogenes). VIRMA, YTHDF1, and YTHDF3 are the most commonly upregulated genes, representing 18%, 19%, and 13% of the altered tumor samples, respectively, and those with the highest upregulated-to-downregulated ratio, together with RBM15 and HAKAI. In contrast, YTHDF2 is the most frequently downregulated factor (5% of tumor samples; **Figure 1D**). Upregulation is also predominant at the level of single tumor types (19/31 tumors examined, paired Wilcox test adjusted p-value < 0.05), with VIRMA, YTHDF1, and YTHDF3 remaining the most upregulated genes (**Figure 1D**). Indeed, VIRMA, YTHDF1, and YTHDF3 are upregulated in at least 10% of samples in 21, 19, and 19 tumors, respectively, out of 31 tumor types. Finally, by analyzing the CNAs of the 15 m^6^A factors, we observed no significant difference between amplification and deep deletion frequencies at the pan-cancer and single tumor type level (pan-cancer: paired Wilcoxon test p-value = 0.5; 31/31 tumors examined: paired Wilcoxon test adjusted p-value > 0.05) (**Figure 1E**). Among all, VIRMA, YTHDF1, and YTHDF3 are the most frequently amplified genes, with 6%, 3%, and 3% of tumor samples altered, respectively (**Figure 1E**).

Taken together, these results show that differential expression, as derived from z-scores, is the most common and relevant type of dysregulation in cancer among the m^6^A factors. YTHDF1-3 and VIRMA are the most altered genes, while, contrary to expectations, the METTL3, ALKBH5, and FTO genes, which are intensely studied in cancer for their direct enzymatic role in m^6^A deposition and removal, present much lower levels of alteration. Specifically, they show a dysregulated expression in 6.2%, 9.6%, and 6.6% of tumor samples, respectively (**Figure 1D**), and a frequency of amplification or deep deletion events not exceeding 1% of tumor samples (**Figure 1E**). These results do not support the hypothesis of a driver role for the "enzymatic" m^6^A genes in the tumorigenesis process, while instead, the cytosolic YTHDF readers and the VIRMA component of the MACOM complex may be under expression-selective pressure in tumor evolution.

### 2. Cross-tumor profile of m^6^A factors displays distinct alteration patterns of YTHDF1, YTHDF2, YTHDF3 and VIRMA

Does the identified dysregulated expression of some of the m^6^A factors suggest that they may play a prominent role in cancer across different tumor types? To address this question, we applied a non-negative matrix factorization (NMF) approach and clustered tumor samples according to their expression dysregulation, obtaining fourteen groups (“classes”) (**Figure 2A**, **Supplementary Table S5**). Although most classes include samples altered in multiple m^6^A factor genes, each class is enriched in only one of them (classes 1, 2, 3, 4, 5, 6, 9, 10, 11, 12, 13, and 14) or two (classes 7 and 8) (**Figure 2B**, **Supplementary Table S6**), the latter case corresponding to genes which lay on the same chromosome arm.

**Figure 2.**
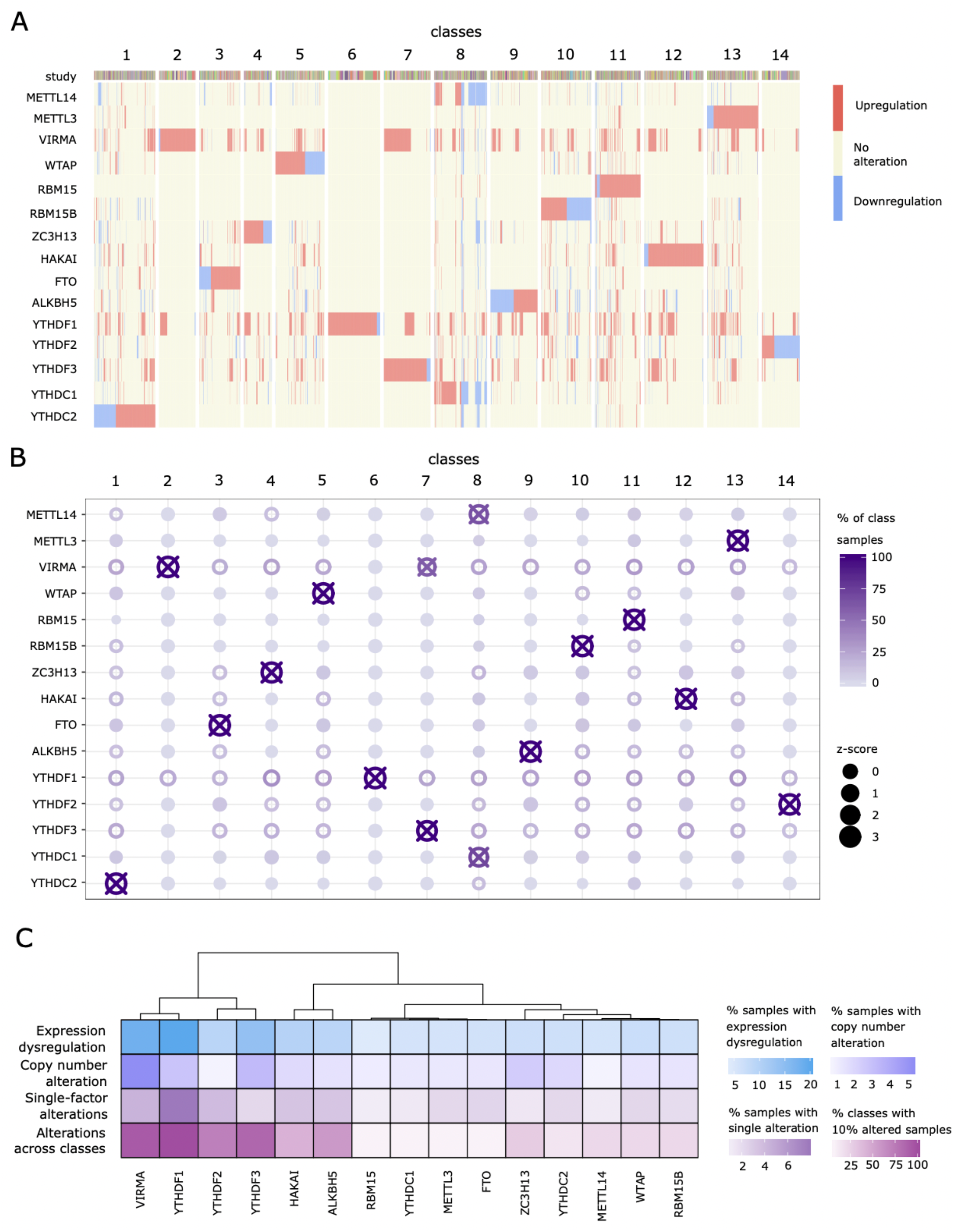
YTHDF1, YTHDF2, YTHDF3, and VIRMA are the most frequently altered genes and the only ones uniquely altered in many tumors. A) The heatmap shows the 14 classes of tumor samples obtained through the NMF clustering method according to their m^6^A factor expression alterations (upregulation: z-score => 2; downregulation: z-score <= -2). Columns correspond to individual samples, subdivided into classes. The tumor type composition is displayed by colors in the upper section. Rows encompass the 15 m^6^A factors under consideration. B) The heatmap displays the frequency of m^6^A factor expression alterations in 14 classes. Color represents the percentage of class samples altered in each m^6^A factor. Size represents the z-scaled percentage of class samples altered in each m^6^A factor. Points with borders indicate m^6^A factors altered in at least 10% of the class samples, and points with crosses indicate m^6^A factors with z-score >= 1. C) Hierarchical clustering based on m^6^A factor expression dysregulations, copy number alterations, and NMF features. The heatmap shows the percentage of tumor samples with expression (up or down-regulations) and copy number (amplification or deep deletions) alterations in each m^6^A factor. The NMF clustering is summarized by displaying the percentage of tumor samples with unique expression alterations in each m^6^A factor as well as the percentage of classes showing at least 10% of samples altered in the corresponding m^6^A factor.

Classes 2, 6, 7, and 14 are the only classes composed of samples with expression alterations exclusively affecting a reduced number (maximum 4) of m^6^A core factors. Interestingly, these are the factors we have previously identified as the most dysregulated in cancer (YTHDF1-3/VIRMA) (**Figure 2A,B**), further suggesting the driving role of these genes in tumorigenesis. While the NMF clustering distributed most tumor types in all classes, almost proportionally to their sample size (**Figure S2B**), classes 2, 6, 7, and 14 are significantly enriched for 5 tumor types (**Supplementary Table S7**). In detail, class 14 is enriched in low-grade glioma (LGG) (18% of class samples) and, to a lesser extent, in pheochromocytoma and paraganglioma (PCPG), while classes 2, 6, and 7 are enriched in urothelial bladder carcinoma (BLCA), stomach adenocarcinoma (STAD) and testicular germ cell tumors (TGCT) samples, respectively (**Supplementary Table S7**).

We subsequently evaluated the expression alteration frequency of the 15 m^6^A factors across the 14 classes and observed that, again, YTHDF1, YTHDF2, YTHDF3, and VIRMA are the most altered genes among the m^6^A core factors (**Figure 2B**). In addition, we also observed that the expression levels of VIRMA in class 2, YTHDF1 in class 6, and VIRMA and YTHDF3 in class 7 are higher than those of the other classes (**Figure S2A**), whereas the expression of YTHDF2 in class 14 is lower than that of the other classes (**Figure S2A**). Overall, these genes are not only uniquely altered in many tumors but also the only ones exhibiting alterations in a cross-class fashion.

Putting together the percentage of tumor samples with unique expression alterations in each m^6^A factor, the percentage of classes with at least 10% of samples altered in the corresponding m^6^A factor, and the percentage of samples with expression dysregulation and CNAs (**Figure 2C**), YTHDF1, YTHDF2, YTHDF3, and VIRMA are clearly the genes to be prioritized for their causative role in oncogenesis and cancer progression.

### 3. YTHDF1, YTHDF2, YTHDF3, and VIRMA expression dysregulation is driven by broad copy number alterations

Following the identification of the most frequently altered genes in the m^6^A core group, we mapped the tumor types where their expression dysregulation primarily occurs (**Figure 3A**, **Supplementary Table S8**). Colon and rectum adenocarcinoma (COAD/READ) and STAD, two gastrointestinal tumors, are enriched in YTHDF1-upregulated samples (adj p-value < 10^-3^), while LGG and PCPG, two neural tumors, are enriched in YTHDF2-downregulated samples (adj p-value < 10^-3^). Similarly, ovarian serous cystadenocarcinoma (OV) and uterine carcinosarcoma (UCS), two gynecological tumors, are enriched in YTHDF3-upregulated samples (adj p-value < 10^-3^ and adj p-value=0.004, respectively), while hepatocellular liver carcinoma (LIHC) is enriched in samples with VIRMA upregulation (adj p-value < 10^-3^). Finally, COAD/READ, TGCT, and uveal melanoma (UVM) are enriched in YTHDF3/VIRMA-upregulated samples (adj p-value < 0.003). We confirmed a significant association between the observed mRNA-level dysregulation and the corresponding protein level for available tissues by exploring protein data present in a recent multi-omics dataset of 2002 primary tumors (54) (**Figure S3A**; **Supplementary Table S9**).

**Figure 3.**
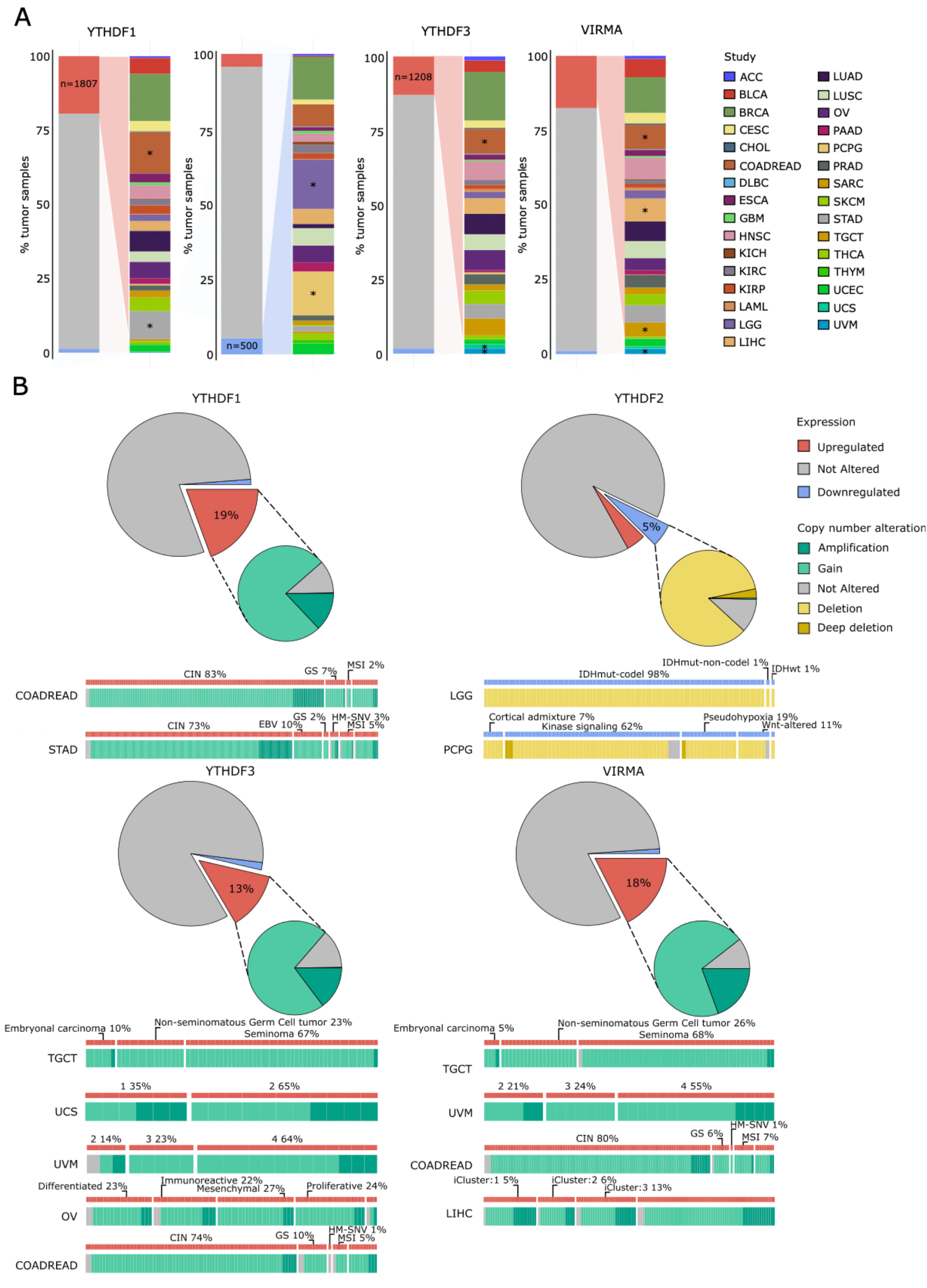
Chromosomal CNAs drive expression dysregulations of key dysregulated m^6^A genes in cancer. A) The plots show the percentage of samples with expression alterations in the most frequently altered m^6^A factors. The number of samples associated with the selected direction of expression dysregulation (upregulation or downregulation) is displayed along with the corresponding percentages of these samples within each tumor type. Enrichment analysis was performed with the Fisher exact test: * adjusted p-value <= 0.05 and odds ratio >= 3. B) Pie charts show the percentage of samples with expression alterations in the most frequently altered m^6^A factors and the corresponding percentage of samples with high/low levels of copy number alterations. The molecular subtypes associated with samples displaying expression dysregulation in these frequently altered m^6^A factors within the enriched tumor types are displayed under the respective pie charts.

To determine if the observed expression dysregulations were due to focused CNAs, chromosomal instability, or altered expression regulations, we examined genomic variations around the YTHDF1, YTHDF2, YTHDF3, and VIRMA loci, considering broad and focal gains as well as heterozygous and homozygous deletions using both low and high-level CNA thresholds. This analysis revealed the widespread involvement of CNAs in mRNA-level dysregulations by showing collinearity between CNAs and expression changes across tumor types (**Figure 3B**). Indeed, in almost all tumor samples enriched for dysregulated YTHDF1-3/VIRMA expression, there are concordant gain/loss events in at least 94% of cases (**Figure 3B**). We next evaluated whether these CNAs were focused or involved broader chromosomal regions and observed widespread gains and losses along the genes of the entire investigated chromosome arms (**Figure S3B**). For instance, 8q gains around VIRMA and YTHDF3 involved at least 80% of genes in 85% of the tumor samples enriched for VIRMA and YTHDF3 upregulated expression. Similarly, tumor samples enriched in YTHDF1 and YTHDF2 alterations show widespread 20q gain and 1p deletion in 94% and 96% of cases, respectively. Since this type of genomic imbalance is characteristic of chromosomal instability (CIN), a well-known and transversal tumor phenotype, we assessed the prevalence of this feature among the tumor samples enriched for dysregulated YTHDF1-3 and VIRMA expression. As expected, we observed a high frequency of samples harboring this specific phenotype (**Figure 3B**). For example, the majority of COAD/READ and STAD altered samples belong to the CIN subtype, characterized by frequent amplification events in 20q (YTHDF1) and 8q (YTHDF3/VIRMA) (70–73). Similarly, 8q amplification is common in OV, UCS, LIHC, and UVM altered samples, the latter mostly belonging to subtypes 3 and 4, characterized by 8q whole-arm gain (74–77). Although the 8q amplification event has not been yet reported in TGCT, the altered samples belong to the seminoma subtype, where a positive correlation of VIRMA and YTHDF3 expression has been found (78, 79). Finally, 1p (YTHDF2) loss is a feature of both LGG and PCPG, with LGG altered samples belonging to the IDHmut-codel subtype (characterized by IDH gene mutation and 1p/19q co-deletion (80)), and PCPG altered samples belonging to the kinase signaling subtype (81).

Besides CNAs involving entire chromosomal arms, we evaluated the putative contribution of single nucleotide polymorphisms (SNPs) to the m^6^A factor expression dysregulations. To this end, we first conducted a heritability analysis to examine the contribution of germline genetic variations assuming YTHDF1-3/VIRMA expression dysregulations as phenotypes (**Figure S4A**) and found significant heritability for all four phenotypes, ranging from 6% to 17% (p-value < 0.05). As cis-associations can contribute to the heritability of gene expression levels and are more likely to have a direct impact on gene expression, we then explored cis-expression quantitative trait loci (cis-eQTLs) using the PancanQTL resource (56). Specifically, we identified 130 cis-eQTLs associations between SNPs and the expression of YTHDF1-3/VIRMA across 11 tumor types (**Supplementary Table S10**). Among them, we observed that most of the cis-eQTLs were associated with increased expression of VIRMA in 8 tumor types (**Supplementary Table S10**). Furthermore, for each of the four analyzed m^6^A factors, we computed linkage disequilibrium (LD) between all the SNPs of the associated cis-eQTLs and found that most of them are in strong LD (R^2^ > 0.8) (**Figure S4B-C**). Interestingly, the tumor types where cis-eQTLs were identified often coincide with those exhibiting dysregulated YTHDF1-3/VIRMA expression and an enrichment for the absence of CNAs (**Figure S4D**).

Taken together, these results indicate that dysregulated expression observed for YTHDF1-3/VIRMA is primarily driven by CNAs involving the entire chromosomal arm, often associated with the CIN phenotype. Moreover, we observed a correlation between SNPs and the expression of m^6^A genes, suggesting that alternative mechanisms also play a role in driving the observed expression dysregulations.

### 4. Evolutionary dependency analysis reveals m^6^A factors implication in cancer driver pathways

Having identified CNAs as the main drivers of expression dysregulation in the most frequently altered m^6^A genes in cancer, we wondered whether alterations in cancer driver genes might also contribute to this phenomenon. We thus explored the potential of trans-interactions between m^6^A factors and cancer driver genes employing the SELECT algorithm, introduced for scoring evolutionary dependencies (59, 82). By applying the SELECT algorithm to the expression dysregulations, amplifications, and deep deletion events, we identified patterns of co-occurrence and mutual exclusivity among the m^6^A factors, as well as between the m^6^A factors and cancer driver genes (n=514 derived from OncoKB). In detail, among the most dysregulated m^6^A factors, we identified significant co-occurrence between VIRMA and YTHDF3 in terms of amplification and up/down-regulations (**Figure 4A**). This co-occurrence explained their observed association in the NMF clustering class 7 (**Figure 2A**) and is likely due to their close proximity on chromosome 8q, which implicates them in frequent pan-cancer 8q gains. In addition, we also observed significant co-occurrence between VIRMA and YTHDC1 downregulations, and between VIRMA and ZC3H13 upregulations. Furthermore, YTHDF3 amplification displays significant mutual exclusivity with ZC3H13 amplification, while YTHDF1 downregulation shows significant mutual exclusivity with YTHDF3 downregulation (**Figure 4A**). The m^6^A factor dysregulations also exhibit significant co-occurrences with expression alterations (574 gene pairs) or CNAs (307 gene pairs) of cancer driver genes (**Supplementary Table S11**). These co-occurrences can be attributed to genomic proximity or may provide insights into shared pathways. Specifically, co-occurrences with CNAs exclusively involve cancer driver genes located on the same chromosomal arms as the considered m^6^A factor (**Supplementary Table S11**), confirming the already established strong influence of genomic proximity on the frequency of concurrent events (82). To exclude hitchhiking effects of the m^6^A factors by the cancer driver genes, we thus investigated the presence of cancer driver genes within a maximum distance of 1 million base pairs (1Mb) from the YTHDF1, YTHDF2, YTHDF3, and VIRMA genes. We identified only the SESN2 tumor suppressor (83), located 450 Kb away from YTHDF2. However, deep deletion events simultaneously affecting SESN2 and YTHDF2 in the corresponding enriched tumor types proved to be very rare, considering YTHDF2-downregulated samples as well as all samples (1.94% and 0.32%, respectively) (**Figure S5A**). Therefore, the significant co-occurrences with CNAs are likely a consequence of frequent chromosomal imbalances that involve all genes on the arm rather than a direct dependence of m^6^A factors on cancer drivers.

**Figure 4.**
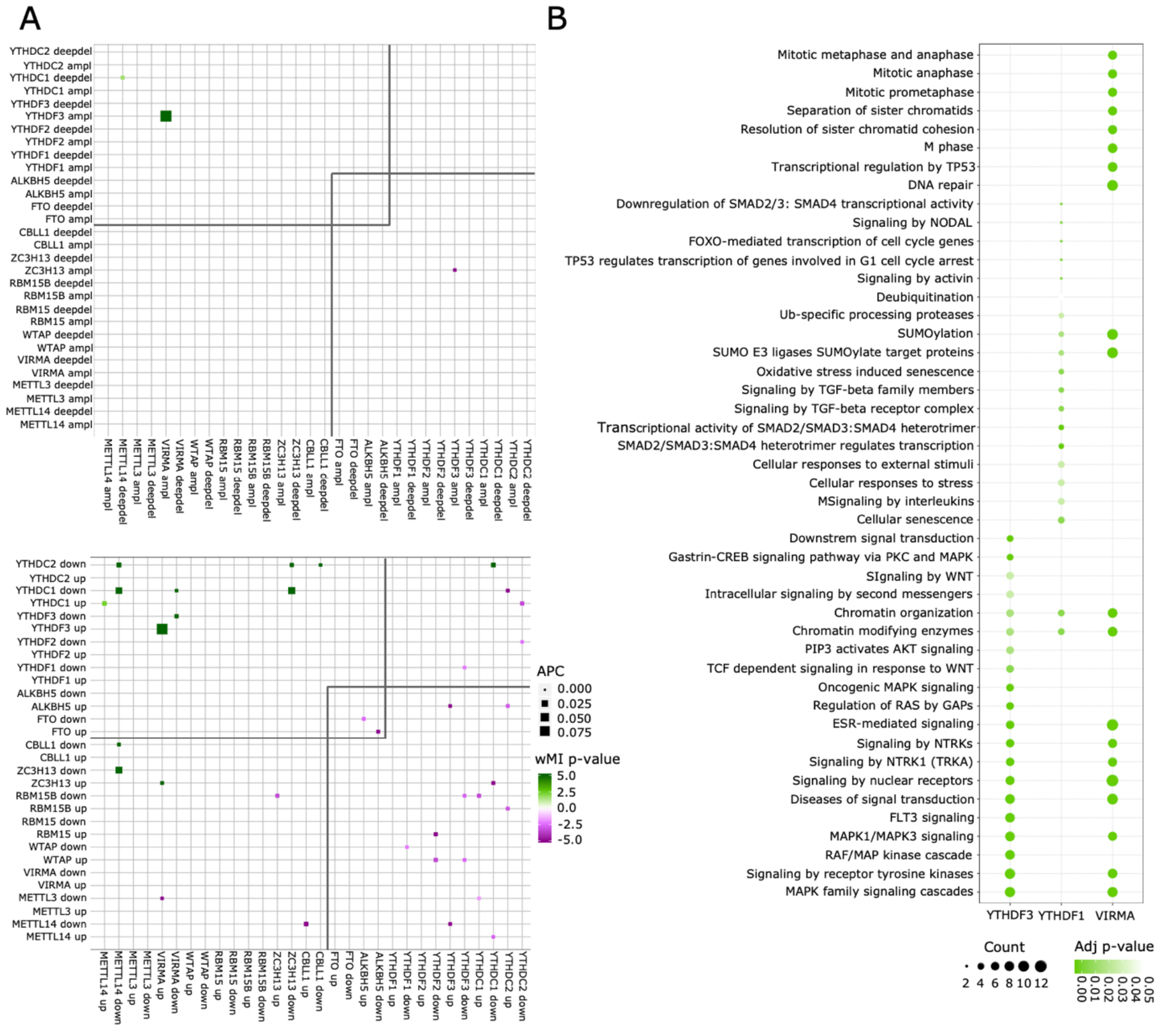
Co-occurrence of m^6^A factors expression dysregulations and m^6^A factors implication in cancer driver pathways. A) The heatmaps show co-occurring (green) and mutually exclusive (violet) copy number (up) and expression (down) alterations of m^6^A factors generated using the SELECT algorithm (59). The color gradient corresponds to the wMI p-values and the size to the effect sizes (ASC-corrected motif score). B) The dot plot shows the results of the pathway enrichment analysis performed on the cancer driver genes co-occurring with expression dysregulations of the most altered m^6^A genes. The size indicates the number of genes overlapping the corresponding pathway terms, and the color indicates the enrichment adjusted p-value (adjusted p-value <= 0.05). Only the first 20 enriched pathway terms for each condition are displayed.

The SELECT algorithm has also been shown to identify significant co-occurring alterations within genes belonging to the same pathways rather than different pathways (82). Consequently, we performed a pathway enrichment analysis on the cancer driver genes that co-occur in terms of expression with the most dysregulated m^6^A factors, specifically focusing on genes located on different chromosomes (**Figure 4B, Supplementary Table S12**). We found that YTHDF2 alterations exclusively co-occur with genes located on the same chromosome. In contrast, YTHDF1 expression dysregulations co-occur with cancer driver genes involved in SUMOylation and TGFβ/SMAD signaling pathways, while VIRMA and YTHDF3 co-occur with genes involved in the MAPK, Wnt, and PI3K/AKT pathways, with VIRMA being associated with SUMOylation as well. In summary, these findings indicate that the most dysregulated genes may contribute to pathways shared with cancer driver genes, and these behaviors may have relevance not only at a tumor-specific level but also across various cancer types.

### 5. Expression dysregulation of the YTHDF1, YTHDF2, YTHDF3, and VIRMA m^6^A factors has pan-cancer predictive prognostic power

To evaluate the prognostic significance of the altered expression of the 15 m^6^A genes in cancer, we conducted a multivariate Cox regression analysis for each gene across all tumor types, assessing increased and decreased expression separately (**Figure 5A**; **Supplementary Table S13**). Interestingly, upregulation of VIRMA, YTHDF1, YTHDF3, RBM15, and RBM15B is associated with a worse prognosis compared to non-upregulated tumor samples (Hazard Ratio (HR)=1.1 [1-1.2], p-value=0.01; HR=1.1 [0.98-1.2], p-value=0.1; HR=1.1 [1-1.3], p-value=0.02; HR=1.2 [0.99-1.4], p-value=0.06; HR=1.2 [0.97-1.4], p-value=0.1, respectively), while downregulation of ALKBH5 is also associated with a worse prognosis (HR=1.2 [0.98-1.4], p-value=0.1). On the contrary, METTL14 upregulation and YTHDF2 downregulation predict a better prognosis (HR=0.79 [0.63-1], p-value=0.06; HR=0.87 [0.73-1], p-value=0.1). To ensure that the cross-tumor survival analyses were unbiased by variations in the median progression-free interval (PFI) times between tumor types, we performed a multivariate Cox regression analysis on quantile normalized PFI times, which showed 87% agreement with the previous analysis (**Supplementary Table S14**). Overall, these data indicate that among the 15 core m^6^A genes, YTHDF1, YTHDF2, YTHDF3, and VIRMA are not only the most altered in expression but also possess pan-cancer prognostic significance.

**Figure 5.**
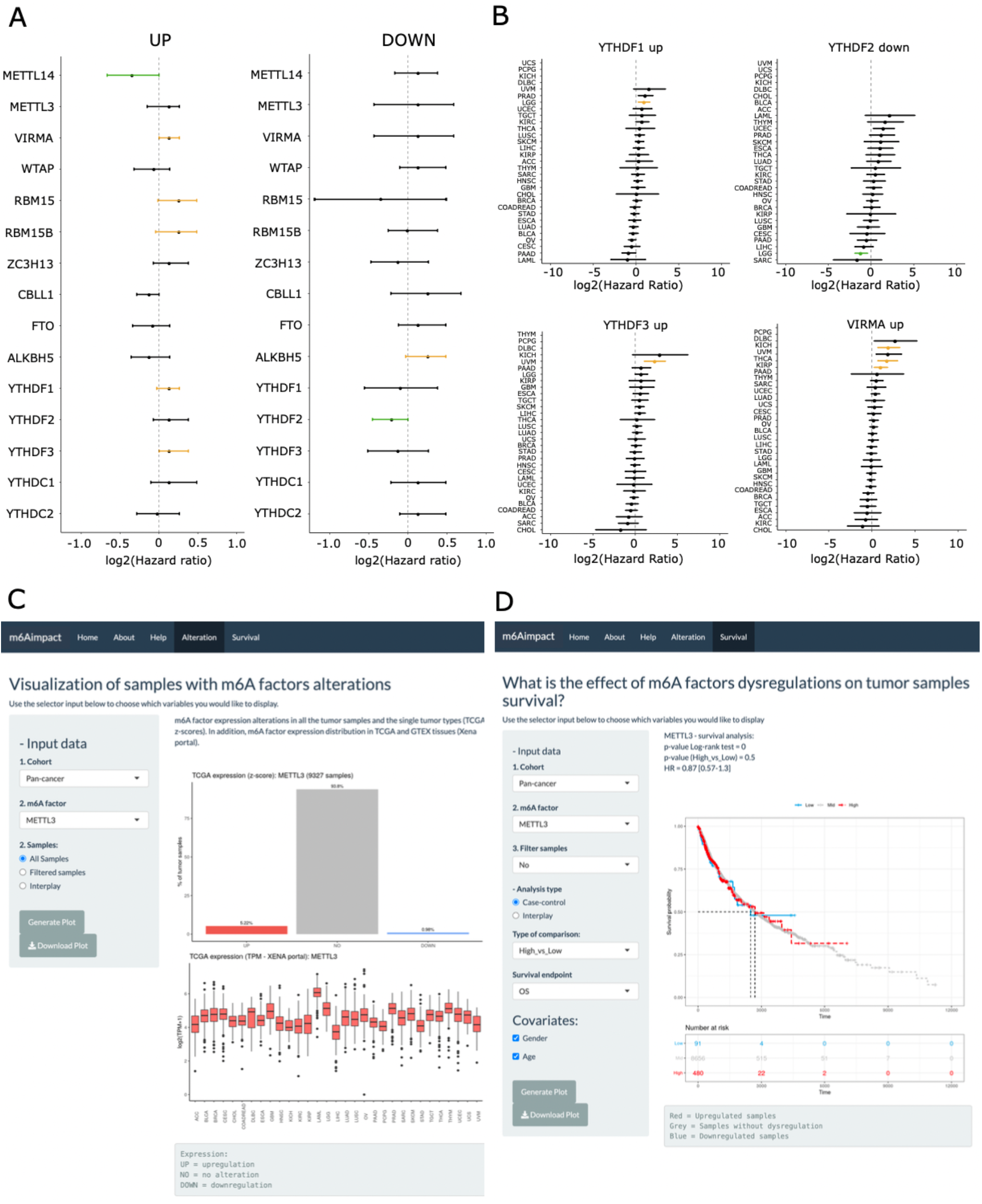
The most frequently dysregulated m^6^A genes show pan-cancer and single-tumor prognostic roles. A) The forest plot shows the results of the multivariate Cox regression analyses performed on samples with m^6^A factors upregulation or downregulation at the pan-cancer level. Yellow/green m^6^A factors are significantly associated with a worse or better prognosis, respectively (p-value < 0.1). B) The forest plots show the results of multivariate Cox regression analyses performed on samples with YTHDF1 upregulation, YTHDF2 downregulation, YHDF3 upregulation, and VIRMA upregulation in individual tumor types. Yellow/green tumor types are significantly associated with a worse/better prognosis (adjusted p-value < 0.1). C) Example search for “Alteration Analysis”. The figure shows the expression distribution of METTL3 at the pan-cancer level and in each single tumor type. D) Example search for “Survival Analysis”. The figure shows the effects of METTL3 expression dysregulation on survival, specifically comparing the impact of upregulation versus downregulation on overall survival (OS) as the endpoint.

Subsequently, we evaluated if the prognostic role of these four genes was consistent across individual tumor types (**Supplementary Table S15**). Our findings indicate that the upregulation of YTHDF3 and VIRMA in UVM samples, as well as the upregulation of VIRMA in kidney renal papillary cell carcinoma (KIRP) and pancreatic adenocarcinoma (PAAD) samples, is associated with a significantly worse prognosis (**Figure 5B**). In contrast, in LGG samples, YTHDF1 upregulation and YTHDF2 downregulation are significantly linked to worse and better prognosis, respectively (**Figure 5B**). When we restricted the survival analysis to samples clustered in the four classes where VIRMA, YTHDF1, YTHDF2 and YTHDF3 are the only altered genes (class 2, 6, 7, 14; **Figure S6A**), we confirmed the results for LGG samples, with YTHDF1 upregulation and YTHDF2 downregulation being associated with worse and better prognosis in class 6 and 14, respectively (**Supplementary Table S16**). However, in class 7, we did not confirm the association between YTHDF3/VIRMA upregulation and unfavorable prognosis in UVM samples but observed it in lung squamous cell carcinoma (LUSC) and OV samples (**Figure S6A**). Overall, despite numerous studies exploring the outcome of various m^6^A factors in individual tumor types, the comprehensive analysis of expression alterations across multiple factors reveals only a few significant associations.

Finally, to allow systematic exploration of the impact of m^6^A factor dysregulations on cancer survival, we developed m6Aimpact, a web application freely accessible a https://ltg.cibio.unitn.it/m6Aimpact/. Users can select among the 15 m^6^A core factors and examine their effect on survival based on expression, CNAs, or both (**Figure 5C, D**). Our web application offers additional features to enhance analysis, including the option to exclude samples with both expression and CNA data. Users can also investigate interplays between pairs of genes, which may include both m^6^A factors and cancer driver genes. These analyses can be conducted at the pan-cancer level or for single tumor types. Furthermore, users can choose from multiple survival endpoints, such as overall survival (OS) or progression-free interval (PFI), and select various covariates to be added to the Cox Proportional Hazard model. m6Aimpact provides an intuitive platform for gaining further insights into the relationship between m^6^A regulators and cancer prognosis.

## Discussion

Over the past decade, the dynamic and reversible m^6^A modification has emerged as a key regulatory mechanism in cancer. However, despite the publication of more than one thousand research reports only in the last four years, the overall significance of m^6^A and the specific roles of its controlling factors in different cancer types remain controversial. Indeed, no clear consensus emerges from the studies regarding the tendency of individual altered m^6^A factors to behave as gain-of-function or loss-of-function actors along the tumorigenic process in a given tissue (84). A comprehensive analysis of the alterations present in the genes encoding the core m^6^A machinery proteins could thus provide an unbiased picture of their potential roles across various cancer types, which might highlight recurrent events. In this context, we investigated the alterations in terms of mutations, differential expressions, and CNAs of 15 m^6^A factors in samples of 31 cancer types from the TCGA dataset. Specifically, we included a gene in the “m^6^A factor” category only when substantial evidence was available for this role, and this role was exclusively related to m^6^A regulation. For instance, we considered as m^6^A readers only the 5 proteins containing the YTH domain, whose only reported function is to recognize N^6^-methylated adenines (85–87) and, instead, excluded other RNA-binding proteins, which have been demonstrated to bind m^6^A marks in a sequence-dependent manner but for which an m^6^A-independent role in cancer has also been demonstrated.

Our pan-cancer profile analysis provides a broad overview of the most frequent alterations occurring in the m^6^A machinery genes across multiple cancer types. This broad perspective reveals common alteration patterns and highlights potential shared therapeutic targets that might be missed in single-cancer studies. Although over the past years other studies have aimed to analyze alterations of m^6^A machinery genes to uncover the context-specific roles of the m^6^A factors within individual cancers or to explore the influence of a single m^6^A factor across different tumors (88–95), these attempts present a number of limitations. For instance, many studies have inferred dysregulation of m^6^A factors by comparing their expression in tumors and normal tissues from available databases, which may have introduced bias due to different tissue origins. In addition, several studies have used common consensus clustering algorithms based on the expression dysregulation of m^6^A factors to identify m^6^A modification patterns, which can be sensitive to initial conditions and parameter choices, potentially leading to variability in results and difficulty in interpretation. To overcome these limitations, we used z-scores derived from diploid samples to determine the expression dysregulation of m^6^A machinery genes. This method assumes a normal distribution of gene expression values and identifies samples with extreme expression values, corresponding to an absolute z-score greater than 2. Although this approach allowed us to identify significantly dysregulated genes in our dataset, it may have excluded more subtle changes in gene expression that may still play a role. In addition, we employed the NMF algorithm, to cluster tumor samples according to their expression dysregulation of m^6^A factors. This approach has previously been used as a clustering method to reveal molecular patterns (96, 97) and offers advantages such as high interpretability and the production of biologically meaningful clusters that are consistent with the non-negative nature of gene expression data.

However, our study may not have captured the full spectrum of changes present in a specific cancer type because our methods focus on identifying extreme expression levels and clear patterns and may have missed more subtle but biologically significant changes in gene expression.

Among the different types of alterations analyzed — expression, mutation, and CNA — we identified differential expression as the most common alteration occurring in the core m^6^A machinery genes. Point mutations occur at a very low frequency across tumors, as well as high-level CNAs, while by including both high- and low-level CNAs, we observed quite abundant alterations in the m^6^A factors. In this context, we showed that differential expression of the m^6^A factors is mainly driven by chromosome arm-level CNAs, as dysregulated samples are often classified in CIN tumor subtypes. GISTIC scores used to identify CNAs are obtained by considering their frequency and amplitude in the samples analyzed. Extreme scores corresponding to amplification and deep deletion events may not have been reported for genes meeting the threshold only in a subset of tumor samples, thus not accurately reflecting the true copy number alteration status of a gene. In addition, the use of values corresponding to gain and deletion events may include genes with limited copy number alterations. The concordant pattern, between expression dysregulation and CNAs, was observed in other studies as well but without identifying alternative mechanisms of regulations of m^6^A factor expression alterations (90). Our study, in contrast, also explored the role of germline genetic variations, and found significant cis-eQTLs in tumor types exhibiting enrichment in samples with expression alterations in the most frequently altered genes and without collinear CNAs. Finally, to explore trans-interactions between m^6^A factors and cancer driver genes contributing to expression dysregulations, we used the SELECT algorithm, which, unlike the common correlation methods assessing relationships between variables, identifies specific patterns and associations between genes, revealing subsets that co-occur or mutually exclude each other across tumor types and therefore may belong to the same rather than different pathways. For instance, we observed the co-occurrence of YTHDF1 upregulation with cancer driver genes involved in the SUMOylation process and the TGFβ/SMAD signaling pathway. Although evidence is limited, YTHDF1-3 has been shown to facilitate TGFβ-induced EMT and invasion in non-small cell lung cancer (98). Similarly, we found co-occurrences of VIRMA and YTHDF3 upregulation with genes associated with the MAPK, Wnt and PI3K/AKT pathways, with VIRMA also linked to SUMOylation. Again, the supporting data for direct involvement of VIRMA and/or YTHDF3 in these pathways is limited, but some associations have been identified. For example, VIRMA has been shown to activate the JNK/MAPK pathway and induce m^6^A-dependent gefitinib resistance in lung adenocarcinoma cells, and YTHDF3 upregulation has been associated with increased gastric cancer cell growth and invasion through activation of the PI3K/AKT pathway and modulation of the immune microenvironment (99, 100). Therefore, our results may provide insight into pathways that may be affected by m^6^A factor alterations and shed light on putative downstream effector genes that are not only tumor-specific but may also be relevant in other tumors.

Through our pan-cancer profile analysis, the most altered genes among the 15 m^6^A factors were identified, representing the main finding of our study. Unexpectedly, the enzymatic factors responsible for both m^6^A deposition and removal, respectively METTL3 and ALKBH5 or FTO, resulted to be rarely altered either at the single tumor level and in a cross-tumor way. Conversely, our study unveiled that the three YTHDF paralogs and the MACOM component VIRMA are the most frequently altered m^6^A factors.

While not directly involved in the deposition or removal of m^6^A, the three YTHDF paralog genes are widely recognized for their critical roles in driving m^6^A-dependent functions. Indeed, the ability of YTHDF proteins to selectively read m^6^A-modified mRNAs allows them to regulate the expression levels of oncogenes or tumor suppressor genes, thereby influencing various cellular functions crucial in cancer (101). For instance, YTHDF1 overexpression has been shown to promote stemness and drug resistance in liver cancer stem cells (102), and YTHDF2 to stabilize oncogenic drivers in glioblastoma (103), while its inhibition in AML selectively compromises tumor initiation and propagation (104). On the other hand, VIRMA and the other components of the MACOM complex have demonstrated their significance in facilitating the localization and binding of the MAC complex to its target mRNAs, consequently influencing its overall profile of activity (4). In support of the crucial role of VIRMA, two independent studies have shown that its silencing leads to a more significant reduction in m^6^A levels compared to the knockout of METTL3/METTL14 (105, 106). Nevertheless, there are still fewer reports of VIRMA involvement in cancer initiation and progression, with respect to the catalytic m^6^A factors.

In addition to identifying the three YTHDF paralogs and VIRMA as the most frequently altered m^6^A factors, our study also showed that these genes are the only m^6^A core factors uniquely altered in samples from different tumor types and that their alteration is associated with patient survival across different cancers. These findings are supported by other recent studies which also reported that these genes are frequently overexpressed across tumor types and are associated with poor prognosis (91, 92). In the context of single tumor survival analysis, while our approach may have missed findings highlighted in the literature for their importance in specific tumor subtypes or using specific survival endpoints, we observed opposite prognostic effects of YTHDF1 and YTHDF2 upregulation and downregulation, respectively, in LGG patients. Recent research has shown that YTHDF2 upregulation is associated with poor prognosis in LGG and is positively correlated with IDH1 expression (107). These findings were also supported by the work of Du et al. and Liu et al., who showed a correlation between YTHDF2 expression and tumor-infiltrating immune cells in LGG (108, 109).

The identification of the YTHDF paralogs and VIRMA as the most frequently altered genes across cancers in our study suggests their central role in tumorigenesis and highlights their potential as promising therapeutic targets. To date, the m^6^A field has mainly focused on METTL3 and FTO, both in terms of studying their role and association with cancer, and at the therapeutic level, due to their enzymatic nature (36, 110). Indeed, a number of small-molecule inhibitors have been proposed to target METTL3, and one of them, STC-15, is now an oral drug in phase I trial in advanced cancers (111). Currently, there is limited research into the druggability of VIRMA and specific inhibitors are lacking. On the other hand, the three YTHDF paralogs, which are all mechanistically implicated in different tumors and have been shown to largely compensate each other (112, 113), have started to be extensively studied from a druggability point of view. Indeed, since their m^6^A RNA recognition pocket is almost identical (114, 115), they could represent a potential cumulative cancer drug target. The first YTHDF inhibitors have been reported. These molecules can either act on all three paralogs simultaneously (116) or be selective for a specific paralog (117–119). In this context, further studies focusing on the role of YTHDF and VIRMA in cancer and their potential inhibition may be crucial to determine their therapeutic viability and importance in cancer treatment.

Globally, our pan-cancer analysis revealed that differential expression is the most prevalent alteration in m^6^A machinery genes, primarily driven by chromosome arm-level CNAs, as dysregulated samples are often classified in CIN tumor subtypes. However, other minor or tumor-specific mechanisms, such as the presence of cis-eQTLs, could also explain the m^6^A factor expression alterations. Among the m^6^A core factors, we found that the three YTHDF paralogs and VIRMA are the most frequently altered genes and are associated with survival prognosis. These results challenge the current focus on METTL3 and FTO, suggesting VIRMA and YTHDFs as promising new targets for cancer therapy. Taken together, these findings provide an innovative perspective on the role of m^6^A genes in cancer and set a potentially valuable basis for future studies aimed at experimentally validating the alterations in these genes in individual tumor types, exploring their druggability or testing their combination as new therapeutic strategies.

## Data availability

The gene expression, copy number alteration, and mutation data of TCGA tumor samples can be found at the cBioPortal (https://www.cbioportal.org/). The gene expression of tumor samples can be found at the Xena Portal (http://xena.ucsc.edu/)(49). Software and resources used for the analyses are described in each method section. All results generated in this study can be found in supplementary tables. The web application is freely available to the public at https://ltg.cibio.unitn.it/m6Aimpact/.

## Supplementary Data

Supplementary Data are available at NAR Online

## Funding

This research was supported by the Fondazione Italiana per la Lotta al Neuroblastoma, by a donation from Enrico and Ivana Zobele, by Fondazione CARITRO (project title: “Terapia differenziante dei tumori: una nuova prospettiva per un classico obiettivo”), and by Associazione Italiana per la Ricerca sul Cancro (AIRC), IG grant 2018 Id. 22075 (A.Q). This work was also supported by the European Union’s Horizon 2020 research and innovation programme under the Marie Skłodowska-Curie grant agreement No 956810 to A.Q., J.M.B., and A.C.; and by the AIRC MFAG grant 2017 Id. 20621 to A.R.; D.S. gratefully acknowledges funding from AIRC (Post-Doc Fellowship, 29868).

## Notes

### Competing Interest Statement

The authors have declared no competing interest.

## References

1. Dominissini, D., Moshitch-Moshkovitz, S., Schwartz, S., Salmon-Divon, M., Ungar, L., Osenberg, S., Cesarkas, K., Jacob-Hirsch, J., Amariglio, N., Kupiec, M., et al. (2012) Topology of the human and mouse m6A RNA methylomes revealed by m6A-seq. Nature, 485, 201–206.

2. Meyer, K.D., Saletore, Y., Zumbo, P., Elemento, O., Mason, C.E. and Jaffrey, S.R. (2012) Comprehensive analysis of mRNA methylation reveals enrichment in 3’ UTRs and near stop codons. Cell, 149, 1635–1646.

3. Zaccara, S., Ries, R.J. and Jaffrey, S.R. (2019) Reading, writing and erasing mRNA methylation. Nat. Rev. Mol. Cell Biol., 20, 608–624.

4. Su, S., Li, S., Deng, T., Gao, M., Yin, Y., Wu, B., Peng, C., Liu, J., Ma, J. and Zhang, K. (2022) Cryo-EM structures of human m6A writer complexes. Cell Res., 32, 982–994.

5. Jia, G., Fu, Y., Zhao, X., Dai, Q., Zheng, G., Yang, Y., Yi, C., Lindahl, T., Pan, T., Yang, Y.-G., et al. (2011) N6-methyladenosine in nuclear RNA is a major substrate of the obesity-associated FTO. Nat. Chem. Biol., 7, 885–887.

6. Zheng, G., Dahl, J.A., Niu, Y., Fedorcsak, P., Huang, C.-M., Li, C.J., Vågbø, C.B., Shi, Y., Wang, W.-L., Song, S.-H., et al. (2013) ALKBH5 is a mammalian RNA demethylase that impacts RNA metabolism and mouse fertility. Mol. Cell, 49, 18–29.

7. Hartmann, A.M., Nayler, O., Schwaiger, F.W., Obermeier, A. and Stamm, S. (1999) The Interaction and Colocalization of Sam68 with the Splicing-associated Factor YT521-B in Nuclear Dots Is Regulated by the Src Family Kinase p59fyn. MBoC, 10, 3909–3926.

8. Wojtas, M., Pandey, R.R., Mendel, M., Homolka, D., Sachidanandam, R. and Pillai, R. (2017) Regulation of m6A transcripts by the 3’→5’ RNA helicase YTHDC2 is essential for a successful meiotic program in the mammalian germline. Mol. Cells, 68, 374–387.e12.

9. Wang, X., Lu, Z., Gomez, A., Hon, G.C., Yue, Y., Han, D., Fu, Y., Parisien, M., Dai, Q., Jia, G., et al. (2014) N6-methyladenosine-dependent regulation of messenger RNA stability. Nature, 505, 117–120.

10. Wang, X., Zhao, B.S., Roundtree, I.A., Lu, Z., Han, D., Ma, H., Weng, X., Chen, K., Shi, H. and He, C. (2015) N(6)-methyladenosine Modulates Messenger RNA Translation Efficiency. Cell, 161, 1388–1399.

11. Li, A., Chen, Y.-S., Ping, X.-L., Yang, X., Xiao, W., Yang, Y., Sun, H.-Y., Zhu, Q., Baidya, P., Wang, X., et al. (2017) Cytoplasmic m6A reader YTHDF3 promotes mRNA translation. Cell Res., 27, 444–447.

12. Shi, H., Wang, X., Lu, Z., Zhao, B.S., Ma, H., Hsu, P.J., Liu, C. and He, C. (2017) YTHDF3 facilitates translation and decay of N6-methyladenosine-modified RNA. Cell Res., 27, 315–328.

13. Edupuganti, R.R., Geiger, S., Lindeboom, R.G.H., Shi, H., Hsu, P.J., Lu, Z., Wang, S.-Y., Baltissen, M.P.A., Jansen, P.W.T.C., Rossa, M., et al. (2017) N6-methyladenosine (m6A) recruits and repels proteins to regulate mRNA homeostasis. Nat. Struct. Mol. Biol., 24, 870–878.

14. Worpenberg, L., Paolantoni, C., Longhi, S., Mulorz, M.M., Lence, T., Wessels, H.-H., Dassi, E., Aiello, G., Sutandy, F.X.R., Scheibe, M., et al. (2021) Ythdf is a N6-methyladenosine reader that modulates Fmr1 target mRNA selection and restricts axonal growth in Drosophila. EMBO J., 40, e104975.

15. Meyer, K.D., Patil, D.P., Zhou, J., Zinoviev, A., Skabkin, M.A., Elemento, O., Pestova, T.V., Qian, S.-B. and Jaffrey, S.R. (2015) 5’ UTR m(6)A Promotes Cap-Independent Translation. Cell, 163, 999–1010.

16. Bell, J.L., Wächter, K., Mühleck, B., Pazaitis, N., Köhn, M., Lederer, M. and Hüttelmaier, S. (2013) Insulin-like growth factor 2 mRNA-binding proteins (IGF2BPs): post-transcriptional drivers of cancer progression? Cell. Mol. Life Sci., 70, 2657–2675.

17. Alarcón, C.R., Goodarzi, H., Lee, H., Liu, X., Tavazoie, S. and Tavazoie, S.F. (2015) HNRNPA2B1 Is a Mediator of m6A-Dependent Nuclear RNA Processing Events. Cell, 162, 1299–1308.

18. Liu, N., Zhou, K.I., Parisien, M., Dai, Q., Diatchenko, L. and Pan, T. (2017) N6-methyladenosine alters RNA structure to regulate binding of a low-complexity protein. Nucleic Acids Res., 45, 6051–6063.

19. Liu, N., Dai, Q., Zheng, G., He, C., Parisien, M. and Pan, T. (2015) N(6)-methyladenosine-dependent RNA structural switches regulate RNA-protein interactions. Nature, 518, 560–564.

20. Flamand, M.N., Tegowski, M. and Meyer, K.D. (2023) The Proteins of mRNA Modification: Writers, Readers, and Erasers. Annu. Rev. Biochem., 92, 145–173.

21. Deng, X., Qing, Y., Horne, D., Huang, H. and Chen, J. (2023) The roles and implications of RNA m6A modification in cancer. Nat. Rev. Clin. Oncol., 20, 507–526.

22. Hong, J., Xu, K. and Lee, J.H. (2022) Biological roles of the RNA m6A modification and its implications in cancer. Exp. Mol. Med., 54, 1822–1832.

23. Niu, X., Yang, Y., Ren, Y., Zhou, S., Mao, Q. and Wang, Y. (2022) Crosstalk between m6A regulators and mRNA during cancer progression. Oncogene, 41, 4407–4419.

24. Li, H., Wu, H., Wang, Q., Ning, S., Xu, S. and Pang, D. (2021) Dual effects of N6-methyladenosine on cancer progression and immunotherapy. Mol. Ther. Nucleic Acids, 24, 25–39.

25. Sun, T., Wu, R. and Ming, L. (2019) The role of m6A RNA methylation in cancer. Biomed. Pharmacother., 112, 108613.

26. Cui, Q., Shi, H., Ye, P., Li, L., Qu, Q., Sun, G., Sun, G., Lu, Z., Huang, Y., Yang, C.-G., et al. (2017) m6A RNA Methylation Regulates the Self-Renewal and Tumorigenesis of Glioblastoma Stem Cells. Cell Rep., 18, 2622–2634.

27. Visvanathan, A., Patil, V., Abdulla, S., Hoheisel, J.D. and Somasundaram, K. (2019) N^6^-Methyladenosine Landscape of Glioma Stem-Like Cells: METTL3 Is Essential for the Expression of Actively Transcribed Genes and Sustenance of the Oncogenic Signaling. Genes, 10.

28. Guo, J., Wu, Y., Du, J., Yang, L., Chen, W., Gong, K., Dai, J., Miao, S., Jin, D. and Xi, S. (2018) Deregulation of UBE2C-mediated autophagy repression aggravates NSCLC progression. Oncogenesis, 7, 49.

29. Chao, Y., Shang, J. and Ji, W. (2020) ALKBH5-m6A-FOXM1 signaling axis promotes proliferation and invasion of lung adenocarcinoma cells under intermittent hypoxia. Biochem. Biophys. Res. Commun., 521, 499–506.

30. Jin, D., Guo, J., Wu, Y., Yang, L., Wang, X., Du, J., Dai, J., Chen, W., Gong, K., Miao, S., et al. (2020) m6A demethylase ALKBH5 inhibits tumor growth and metastasis by reducing YTHDFs-mediated YAP expression and inhibiting miR-107/LATS2-mediated YAP activity in NSCLC. Mol. Cancer, 19, 40.

31. Vu, L.P., Pickering, B.F., Cheng, Y., Zaccara, S., Nguyen, D., Minuesa, G., Chou, T., Chow, A., Saletore, Y., MacKay, M., et al. (2017) The N6-methyladenosine (m6A)-forming enzyme METTL3 controls myeloid differentiation of normal hematopoietic and leukemia cells. Nat. Med., 23, 1369–1376.

32. Barbieri, I., Tzelepis, K., Pandolfini, L., Shi, J., Millán-Zambrano, G., Robson, S.C., Aspris, D., Migliori, V., Bannister, A.J., Han, N., et al. (2017) Promoter-bound METTL3 maintains myeloid leukaemia by m6A-dependent translation control. Nature, 552, 126–131.

33. Yankova, E., Blackaby, W., Albertella, M., Rak, J., De Braekeleer, E., Tsagkogeorga, G., Pilka, E.S., Aspris, D., Leggate, D., Hendrick, A.G., et al. (2021) Small-molecule inhibition of METTL3 as a strategy against myeloid leukaemia. Nature, 593, 597–601.

34. Li, Z., Weng, H., Su, R., Weng, X., Zuo, Z., Li, C., Huang, H., Nachtergaele, S., Dong, L., Hu, C., et al. (2017) FTO Plays an Oncogenic Role in Acute Myeloid Leukemia as a N6-Methyladenosine RNA Demethylase. Cancer Cell, 31, 127–141.

35. Su, R., Dong, L., Li, Y., Gao, M., Han, L., Wunderlich, M., Deng, X., Li, H., Huang, Y., Gao, L., et al. (2020) Targeting FTO Suppresses Cancer Stem Cell Maintenance and Immune Evasion. Cancer Cell, 38, 79–96.e11.

36. Huang, Y., Su, R., Sheng, Y., Dong, L., Dong, Z., Xu, H., Ni, T., Zhang, Z.S., Zhang, T., Li, C., et al. (2019) Small-Molecule Targeting of Oncogenic FTO Demethylase in Acute Myeloid Leukemia. Cancer Cell, 35, 677–691.e10.

37. Xiao, P., Duan, Z., Liu, Z., Chen, L., Zhang, D., Liu, L., Zhou, C., Gan, J., Dong, Z. and Yang, C.-G. (2023) Rational Design of RNA Demethylase FTO Inhibitors with Enhanced Antileukemia Drug-Like Properties. J. Med. Chem., 66, 9731–9752.

38. Cheng, M., Sheng, L., Gao, Q., Xiong, Q., Zhang, H., Wu, M., Liang, Y., Zhu, F., Zhang, Y., Zhang, X., et al. (2019) The m6A methyltransferase METTL3 promotes bladder cancer progression via AFF4/NF-κB/MYC signaling network. Oncogene, 38, 3667–3680.

39. Tao, L., Mu, X., Chen, H., Jin, D., Zhang, R., Zhao, Y., Fan, J., Cao, M. and Zhou, Z. (2021) FTO modifies the m6A level of MALAT and promotes bladder cancer progression. Clin. Transl. Med., 11, e310.

40. Tan, Z., Shi, S., Xu, J., Liu, X., Lei, Y., Zhang, B., Hua, J., Meng, Q., Wang, W., Yu, X., et al. (2022) RNA N6-methyladenosine demethylase FTO promotes pancreatic cancer progression by inducing the autocrine activity of PDGFC in an m6A-YTHDF2-dependent manner. Oncogene, 41, 2860–2872.

41. Lin, C., Li, T., Wang, Y., Lai, S., Huang, Y., Guo, Z., Zhang, X. and Weng, S. (2023) METTL3 enhances pancreatic ductal adenocarcinoma progression and gemcitabine resistance through modifying DDX23 mRNA N6 adenosine methylation. Cell Death Dis., 14, 221.

42. Li, J., Han, Y., Zhang, H., Qian, Z., Jia, W., Gao, Y., Zheng, H. and Li, B. (2019) The m6A demethylase FTO promotes the growth of lung cancer cells by regulating the m6A level of USP7 mRNA. Biochem. Biophys. Res. Commun., 512, 479–485.

43. Jin, D., Guo, J., Wu, Y., Du, J., Yang, L., Wang, X., Di, W., Hu, B., An, J., Kong, L., et al. (2021) m6A mRNA methylation initiated by METTL3 directly promotes YAP translation and increases YAP activity by regulating the MALAT1-miR-1914-3p-YAP axis to induce NSCLC drug resistance and metastasis. J. Hematol. Oncol., 14, 32.

44. Dou, X., Wang, Z., Lu, W., Miao, L. and Zhao, Y. (2022) METTL3 promotes non-small cell lung cancer (NSCLC) cell proliferation and colony formation in a m6A-YTHDF1 dependent way. BMC Pulm. Med., 22, 324.

45. Cerami, E., Gao, J., Dogrusoz, U., Gross, B.E., Sumer, S.O., Aksoy, B.A., Jacobsen, A., Byrne, C.J., Heuer, M.L., Larsson, E., et al. (2012) The cBio cancer genomics portal: an open platform for exploring multidimensional cancer genomics data. Cancer Discov., 2, 401–404.

46. Gao, J., Aksoy, B.A., Dogrusoz, U., Dresdner, G., Gross, B., Sumer, S.O., Sun, Y., Jacobsen, A., Sinha, R., Larsson, E., et al. (2013) Integrative analysis of complex cancer genomics and clinical profiles using the cBioPortal. Sci. Signal., 6, l1.

47. Mermel, C.H., Schumacher, S.E., Hill, B., Meyerson, M.L., Beroukhim, R. and Getz, G. (2011) GISTIC2.0 facilitates sensitive and confident localization of the targets of focal somatic copy-number alteration in human cancers. Genome Biology, 12.

48. Colaprico, A., Silva, T.C., Olsen, C., Garofano, L., Cava, C., Garolini, D., Sabedot, T.S., Malta, T.M., Pagnotta, S.M., Castiglioni, I., et al. (2016) TCGAbiolinks: an R/Bioconductor package for integrative analysis of TCGA data. Nucleic Acids Res., 44, e71.

49. Goldman, M.J., Craft, B., Hastie, M., Repečka, K., McDade, F., Kamath, A., Banerjee, A., Luo, Y., Rogers, D., Brooks, A.N., et al. (2020) Visualizing and interpreting cancer genomics data via the Xena platform. Nat. Biotechnol., 38, 675–678.

50. Gaujoux, R. and Seoighe, C. (2010) A flexible R package for nonnegative matrix factorization. BMC Bioinformatics, 11.

51. Brunet, J.-P., Tamayo, P., Golub, T.R. and Mesirov, J.P. (2004) Metagenes and molecular pattern discovery using matrix factorization. Proc. Natl. Acad. Sci. U. S. A., 101, 4164–4169.

52. Yu, G., Wang, L.-G., Han, Y. and He, Q.-Y. (2012) clusterProfiler: an R package for comparing biological themes among gene clusters. OMICS, 16, 284–287.

53. Yu, G. and He, Q.-Y. (2016) ReactomePA: an R/Bioconductor package for reactome pathway analysis and visualization. Mol. Biosyst., 12, 477–479.

54. Zhang, Y., Chen, F., Chandrashekar, D.S., Varambally, S. and Creighton, C.J. (2022) Proteogenomic characterization of 2002 human cancers reveals pan-cancer molecular subtypes and associated pathways. Nat. Commun., 13, 2669.

55. Yang, J., Benyamin, B., McEvoy, B.P., Gordon, S., Henders, A.K., Nyholt, D.R., Madden, P.A., Heath, A.C., Martin, N.G., Montgomery, G.W., et al. (2010) Common SNPs explain a large proportion of the heritability for human height. Nat. Genet., 42, 565–569.

56. Gong, J., Mei, S., Liu, C., Xiang, Y., Ye, Y., Zhang, Z., Feng, J., Liu, R., Diao, L., Guo, A.-Y., et al. (2018) PancanQTL: systematic identification of cis-eQTLs and trans-eQTLs in 33 cancer types. Nucleic Acids Res., 46, D971–D976.

57. Gerard, D. (2021) Pairwise linkage disequilibrium estimation for polyploids. Mol. Ecol. Resour., 21, 1230–1242.

58. Gu, Z., Gu, L., Eils, R., Schlesner, M. and Brors, B. (2014) circlize Implements and enhances circular visualization in R. Bioinformatics, 30, 2811–2812.

59. Mina, M., Iyer, A., Tavernari, D., Raynaud, F. and Ciriello, G. (2020) Discovering functional evolutionary dependencies in human cancers. Nat. Genet., 52, 1198–1207.

60. Liu, J., Lichtenberg, T., Hoadley, K.A., Poisson, L.M., Lazar, A.J., Cherniack, A.D., Kovatich, A.J., Benz, C.C., Levine, D.A., Lee, A.V., et al. (2018) An Integrated TCGA Pan-Cancer Clinical Data Resource to Drive High-Quality Survival Outcome Analytics. Cell, 173, 400–416.e11.

61. Therneau, T.M. and Foundation, M. (1999) A package for survival analysis in S.

62. Kassambara, A., Kosinski, M., Biecek, P. and Fabian, S. survminer: Drawing Survival Curves using ‘ggplot2’. R package version 0.3.

63. Beroukhim, R., Mermel, C.H., Porter, D., Wei, G., Raychaudhuri, S., Donovan, J., Barretina, J., Boehm, J.S., Dobson, J., Urashima, M., et al. (2010) The landscape of somatic copy-number alteration across human cancers. Nature, 463, 899–905.

64. Pan-cancer analysis of whole genomes (2020) Nature, 578, 82–93.

65. Kumar, S., Warrell, J., Li, S., McGillivray, P.D., Meyerson, W., Salichos, L., Harmanci, A., Martinez-Fundichely, A., Chan, C.W.Y., Nielsen, M.M., et al. (2020) Passenger Mutations in More Than 2, 500 Cancer Genomes: Overall Molecular Functional Impact and Consequences. Cell, 180, 915–927.e16.

66. Martínez-Jiménez, F., Muiños, F., Sentís, I., Deu-Pons, J., Reyes-Salazar, I., Arnedo-Pac, C., Mularoni, L., Pich, O., Bonet, J., Kranas, H., et al. (2020) A compendium of mutational cancer driver genes. Nat. Rev. Cancer, 20, 555–572.

67. Bailey, M.H., Tokheim, C., Porta-Pardo, E., Sengupta, S., Bertrand, D., Weerasinghe, A., Colaprico, A., Wendl, M.C., Kim, J., Reardon, B., et al. (2018) Comprehensive Characterization of Cancer Driver Genes and Mutations. Cell, 174, 1034–1035.

68. Hsiao, H.-H., Yang, M.-Y., Liu, Y.-C., Hsiao, H.-P., Tseng, S.-B., Chao, M.-C., Liu, T.-C. and Lin, S.-F. (2005) RBM15-MKL1 (OTT-MAL) fusion transcript in an adult acute myeloid leukemia patient. Am. J. Hematol., 79, 43–45.

69. Ma, Z., Morris, S.W., Valentine, V., Li, M., Herbrick, J.A., Cui, X., Bouman, D., Li, Y., Mehta, P.K., Nizetic, D., et al. (2001) Fusion of two novel genes, RBM15 and MKL1, in the t(1;22)(p13;q13) of acute megakaryoblastic leukemia. Nat. Genet., 28, 220–221.

70. Buffart, T.E., van Grieken, N.C.T., Tijssen, M., Coffa, J., Ylstra, B., Grabsch, H.I., van de Velde, C.J.H., Carvalho, B. and Meijer, G.A. (2009) High resolution analysis of DNA copy-number aberrations of chromosomes 8, 13, and 20 in gastric cancers. Virchows Arch., 455, 213–223.

71. Zhang, B., Yao, K., Zhou, E., Zhang, L. and Cheng, C. (2021) Chr20q Amplification Defines a Distinct Molecular Subtype of Microsatellite Stable Colorectal Cancer. Cancer Res., 81, 1977–1987.

72. Tan, E.S., Knepper, T.C., Wang, X., Permuth, J.B., Wang, L., Fleming, J.B. and Xie, H. (2022) Copy Number Alterations as Novel Biomarkers and Therapeutic Targets in Colorectal Cancer. Cancers, 14.

73. Liu, Y., Sethi, N.S., Hinoue, T., Schneider, B.G., Cherniack, A.D., Sanchez-Vega, F., Seoane, J.A., Farshidfar, F., Bowlby, R., Islam, M., et al. (2018) Comparative Molecular Analysis of Gastrointestinal Adenocarcinomas. Cancer Cell, 33, 721–735.e8.

74. Cherniack, A.D., Shen, H., Walter, V., Stewart, C., Murray, B.A., Bowlby, R., Hu, X., Ling, S., Soslow, R.A., Broaddus, R.R., et al. (2017) Integrated Molecular Characterization of Uterine Carcinosarcoma. Cancer Cell, 31, 411–423.

75. Cancer Genome Atlas Research Network. Electronic address: and Cancer Genome Atlas Research Network (2017) Comprehensive and Integrative Genomic Characterization of Hepatocellular Carcinoma. Cell, 169, 1327–1341.e23.

76. Network, C.G.A.R. and Others (2011) Integrated genomic analyses of ovarian carcinoma. Nature, 474, 609.

77. Robertson, A.G., Shih, J., Yau, C., Gibb, E.A., Oba, J., Mungall, K.L., Hess, J.M., Uzunangelov, V., Walter, V., Danilova, L., et al. (2018) Integrative Analysis Identifies Four Molecular and Clinical Subsets in Uveal Melanoma. Cancer Cell, 33, 151.

78. Lobo, J., Costa, A.L., Cantante, M., Guimarães, R., Lopes, P., Antunes, L., Braga, I., Oliveira, J., Pelizzola, M., Henrique, R., et al. (2019) m6A RNA modification and its writer/reader VIRMA/YTHDF3 in testicular germ cell tumors: a role in seminoma phenotype maintenance. J. Transl. Med., 17, 79.

79. Shen, H., Shih, J., Hollern, D.P., Wang, L., Bowlby, R., Tickoo, S.K., Thorsson, V., Mungall, A.J., Newton, Y., Hegde, A.M., et al. (2018) Integrated Molecular Characterization of Testicular Germ Cell Tumors. Cell Rep., 23, 3392–3406.

80. Ceccarelli, M., Barthel, F.P., Malta, T.M., Sabedot, T.S., Salama, S.R., Murray, B.A., Morozova, O., Newton, Y., Radenbaugh, A., Pagnotta, S.M., et al. (2016) Molecular Profiling Reveals Biologically Discrete Subsets and Pathways of Progression in Diffuse Glioma. Cell, 164, 550–563.

81. Fishbein, L., Leshchiner, I., Walter, V., Danilova, L., Robertson, A.G., Johnson, A.R., Lichtenberg, T.M., Murray, B.A., Ghayee, H.K., Else, T., et al. (2017) Comprehensive Molecular Characterization of Pheochromocytoma and Paraganglioma. Cancer Cell, 31, 181–193.

82. Mina, M., Raynaud, F., Tavernari, D., Battistello, E., Sungalee, S., Saghafinia, S., Laessle, T., Sanchez-Vega, F., Schultz, N., Oricchio, E., et al. (2017) Conditional Selection of Genomic Alterations Dictates Cancer Evolution and Oncogenic Dependencies. Cancer Cell, 32, 155–168.e6.

83. Ala, M. (2022) Sestrin2 in cancer: a foe or a friend? Biomark. Res., 10, 29.

84. An, Y. and Duan, H. (2022) The role of m6A RNA methylation in cancer metabolism. Mol. Cancer, 21, 14.

85. Luo, S. and Tong, L. (2014) Molecular basis for the recognition of methylated adenines in RNA by the eukaryotic YTH domain. Proc. Natl. Acad. Sci. U. S. A., 111, 13834–13839.

86. Theler, D., Dominguez, C., Blatter, M., Boudet, J. and Allain, F.H.-T. (2014) Solution structure of the YTH domain in complex with N6-methyladenosine RNA: a reader of methylated RNA. Nucleic Acids Res., 42, 13911–13919.

87. Xu, C., Liu, K., Ahmed, H., Loppnau, P., Schapira, M. and Min, J. (2015) Structural Basis for the Discriminative Recognition of N6-Methyladenosine RNA by the Human YT521-B Homology Domain Family of Proteins. J. Biol. Chem., 290, 24902–24913.

88. Shen, S., Zhang, R., Jiang, Y., Li, Y., Lin, L., Liu, Z., Zhao, Y., Shen, H., Hu, Z., Wei, Y., et al. (2021) Comprehensive analyses of m6A regulators and interactive coding and non-coding RNAs across 32 cancer types. Mol. Cancer, 20, 67.

89. Begik, O., Lucas, M.C., Liu, H., Ramirez, J.M., Mattick, J.S. and Novoa, E.M. (2020) Integrative analyses of the RNA modification machinery reveal tissue- and cancer-specific signatures. Genome Biol., 21, 97.

90. Li, Y., Xiao, J., Bai, J., Tian, Y., Qu, Y., Chen, X., Wang, Q., Li, X., Zhang, Y. and Xu, J. (2019) Molecular characterization and clinical relevance of m6A regulators across 33 cancer types. Mol. Cancer, 18, 137.

91. Li, L., Tang, C., Ye, J., Xu, D., Chu, C., Wang, L., Zhou, Q., Gan, S. and Liu, B. (2023) Bioinformatic analysis of m6A ‘reader’ YTH family in pan-cancer as a clinical prognosis biomarker. Sci. Rep., 13, 17350.

92. Ma, C., Zheng, Q., Wang, Y., Li, G., Zhao, M. and Sun, Z. (2023) Pan-cancer analysis and experimental validation revealed the m6A methyltransferase KIAA1429 as a potential biomarker for diagnosis, prognosis, and immunotherapy. Aging, 15, 8664–8691.

93. Zhu, Y., Li, J., Yang, H., Yang, X., Zhang, Y., Yu, X., Li, Y., Chen, G. and Yang, Z. (2023) The potential role of m6A reader YTHDF1 as diagnostic biomarker and the signaling pathways in tumorigenesis and metastasis in pan-cancer. Cell Death Discov, 9, 34.

94. Li, D., Chen, T. and Li, Q.-G. (2023) Identification of a m6A-related ferroptosis signature as a potential predictive biomarker for lung adenocarcinoma. BMC Pulm. Med., 23, 128.

95. Ji, J., Liu, S., Liang, Y. and Zheng, G. (2023) Comprehensive analysis of m6A regulators and relationship with tumor microenvironment, immunotherapy strategies in colorectal adenocarcinoma. BMC Genom Data, 24, 44.

96. Lee, D.D. and Seung, H.S. (1999) Learning the parts of objects by non-negative matrix factorization. Nature, 401, 788–791.

97. Brunet, J.-P., Tamayo, P., Golub, T.R. and Mesirov, J.P. (2004) Metagenes and molecular pattern discovery using matrix factorization. Proc. Natl. Acad. Sci. U. S. A., 101, 4164–4169.

98. Sun, Z., Su, Z., Zhou, Z., Wang, S., Wang, Z., Tong, X., Li, C., Wang, Y., Chen, X., Lei, Z., et al. (2022) RNA demethylase ALKBH5 inhibits TGF-β-induced EMT by regulating TGF-β/SMAD signaling in non-small cell lung cancer. FASEB J., 36, e22283.

99. Lin, X., Ye, R., Li, Z., Zhang, B., Huang, Y., Du, J., Wang, B., Meng, H., Xian, H., Yang, X., et al. (2023) KIAA1429 promotes tumorigenesis and gefitinib resistance in lung adenocarcinoma by activating the JNK/ MAPK pathway in an m6A-dependent manner. Drug Resist. Updat., 66, 100908.

100. Yu, Y., Meng, L.-L., Chen, X.-Y., Fan, H.-N., Chen, M., Zhang, J. and Zhu, J.-S. (2023) m6A reader YTHDF3 is associated with clinical prognosis, related RNA signatures and immunosuppression in gastric cancer. Cell. Signal., 108, 110699.

101. Liao, J., Wei, Y., Liang, J., Wen, J., Chen, X., Zhang, B. and Chu, L. (2022) Insight into the structure, physiological function, and role in cancer of m6A readers—YTH domain-containing proteins. Cell Death Discovery, 8, 1–14.

102. Zhang, X., Su, T., Wu, Y., Cai, Y., Wang, L., Liang, C., Zhou, L., Wang, S., Li, X.-X., Peng, S., et al. (2024) N6-Methyladenosine Reader YTHDF1 Promotes Stemness and Therapeutic Resistance in Hepatocellular Carcinoma by Enhancing NOTCH1 Expression. Cancer Res., 84, 827–840.

103. Dixit, D., Prager, B.C., Gimple, R.C., Poh, H.X., Wang, Y., Wu, Q., Qiu, Z., Kidwell, R.L., Kim, L.J.Y., Xie, Q., et al. (2021) The RNA m6A Reader YTHDF2 Maintains Oncogene Expression and Is a Targetable Dependency in Glioblastoma Stem Cells. Cancer Discov., 11, 480–499.

104. Paris, J., Morgan, M., Campos, J., Spencer, G.J., Shmakova, A., Ivanova, I., Mapperley, C., Lawson, H., Wotherspoon, D.A., Sepulveda, C., et al. (2019) Targeting the RNA m6A Reader YTHDF2 Selectively Compromises Cancer Stem Cells in Acute Myeloid Leukemia. Cell Stem Cell, 25, 137–148.e6.

105. Schwartz, S., Mumbach, M.R., Jovanovic, M., Wang, T., Maciag, K., Bushkin, G.G., Mertins, P., Ter-Ovanesyan, D., Habib, N., Cacchiarelli, D., et al. (2014) Perturbation of m6A writers reveals two distinct classes of mRNA methylation at internal and 5’ sites. Cell Rep., 8, 284–296.

106. Yue, Y., Liu, J., Cui, X., Cao, J., Luo, G., Zhang, Z., Cheng, T., Gao, M., Shu, X., Ma, H., et al. (2018) VIRMA mediates preferential m6A mRNA methylation in 3′UTR and near stop codon and associates with alternative polyadenylation. Cell Discovery, 4, 1–17.

107. Lin, X., Wang, Z., Yang, G., Wen, G. and Zhang, H. (2020) YTHDF2 correlates with tumor immune infiltrates in lower-grade glioma. Aging, 12, 18476–18500.

108. Liu, W., Liu, C., You, J., Chen, Z., Qian, C., Lin, W., Yu, L., Ye, L., Zhao, L. and Zhou, R. (2022) Pan-cancer analysis identifies YTHDF2 as an immunotherapeutic and prognostic biomarker. Front Cell Dev Biol, 10, 954214.

109. Du, J., Ji, H., Ma, S., Jin, J., Mi, S., Hou, K., Dong, J., Wang, F., Zhang, C., Li, Y., et al. (2021) m6A regulator-mediated methylation modification patterns and characteristics of immunity and stemness in low-grade glioma. Brief. Bioinform., 22.

110. Yankova, E., Blackaby, W., Albertella, M., Rak, J., De Braekeleer, E., Tsagkogeorga, G., Pilka, E.S., Aspris, D., Leggate, D., Hendrick, A.G., et al. (2021) Small-molecule inhibition of METTL3 as a strategy against myeloid leukaemia. Nature, 593, 597–601.

111. Blackaby, W.P., Hardick, D.J., Thomas, E.J., Brookfield, F.A., Shepherd, J., Bubert, C. and Ridgill, M.P. (2021) Polyheterocyclic compounds as mettl3 inhibitors. World Patent.

112. Lasman, L., Krupalnik, V., Viukov, S., Mor, N., Aguilera-Castrejon, A., Schneir, D., Bayerl, J., Mizrahi, O., Peles, S., Tawil, S., et al. (2020) Context-dependent functional compensation between Ythdf m6A reader proteins. Genes Dev., 34, 1373–1391.

113. Zaccara, S. and Jaffrey, S.R. (2020) A Unified Model for the Function of YTHDF Proteins in Regulating m6A-Modified mRNA. Cell, 181, 1582–1595.e18.

114. Nai, F., Nachawati, R., Zálešák, F., Wang, X., Li, Y. and Caflisch, A. (2022) Fragment Ligands of the m6A-RNA Reader YTHDF2. ACS Med. Chem. Lett., 13, 1500–1509.

115. Cazzanelli, G., Dalle Vedove, A., Spagnolli, G., Terruzzi, L., Colasurdo, E., Boldrini, A., Patsilinakos, A., Sturlese, M., Grottesi, A., Biasini, E., et al. (2024) Pliability in the m6A-Binding Region Extends Druggability of YTH Domains. J. Chem. Inf. Model., 64, 1682–1690.

116. Micaelli, M., Dalle Vedove, A., Cerofolini, L., Vigna, J., Sighel, D., Zaccara, S., Bonomo, I., Poulentzas, G., Rosatti, E.F., Cazzanelli, G., et al. (2022) Small-Molecule Ebselen Binds to YTHDF Proteins Interfering with the Recognition of N 6-Methyladenosine-Modified RNAs. ACS Pharmacol Transl Sci, 5, 872–891.

117. Wang, L., Dou, X., Chen, S., Yu, X., Huang, X., Zhang, L., Chen, Y., Wang, J., Yang, K., Bugno, J., et al. (2023) YTHDF2 inhibition potentiates radiotherapy antitumor efficacy. Cancer Cell, 41, 1294–1308.e8.

118. Zou, Z., Wei, J., Chen, Y., Kang, Y., Shi, H., Yang, F., Shi, Z., Chen, S., Zhou, Y., Sepich-Poore, C., et al. (2023) FMRP phosphorylation modulates neuronal translation through YTHDF1. Mol. Cell, 83, 4304–4317.e8.

119. Hong, Y.-G., Yang, Z., Chen, Y., Liu, T., Zheng, Y., Zhou, C., Wu, G.-C., Chen, Y., Xia, J., Wen, R., et al. (2023) The RNA m6A Reader YTHDF1 Is Required for Acute Myeloid Leukemia Progression. Cancer Res., 83, 845–860.

