## Supplementary Figures for "The three YTHDF paralogs and VIRMA are the major tumor drivers among the m^6^A core genes in a pan-cancer analysis"

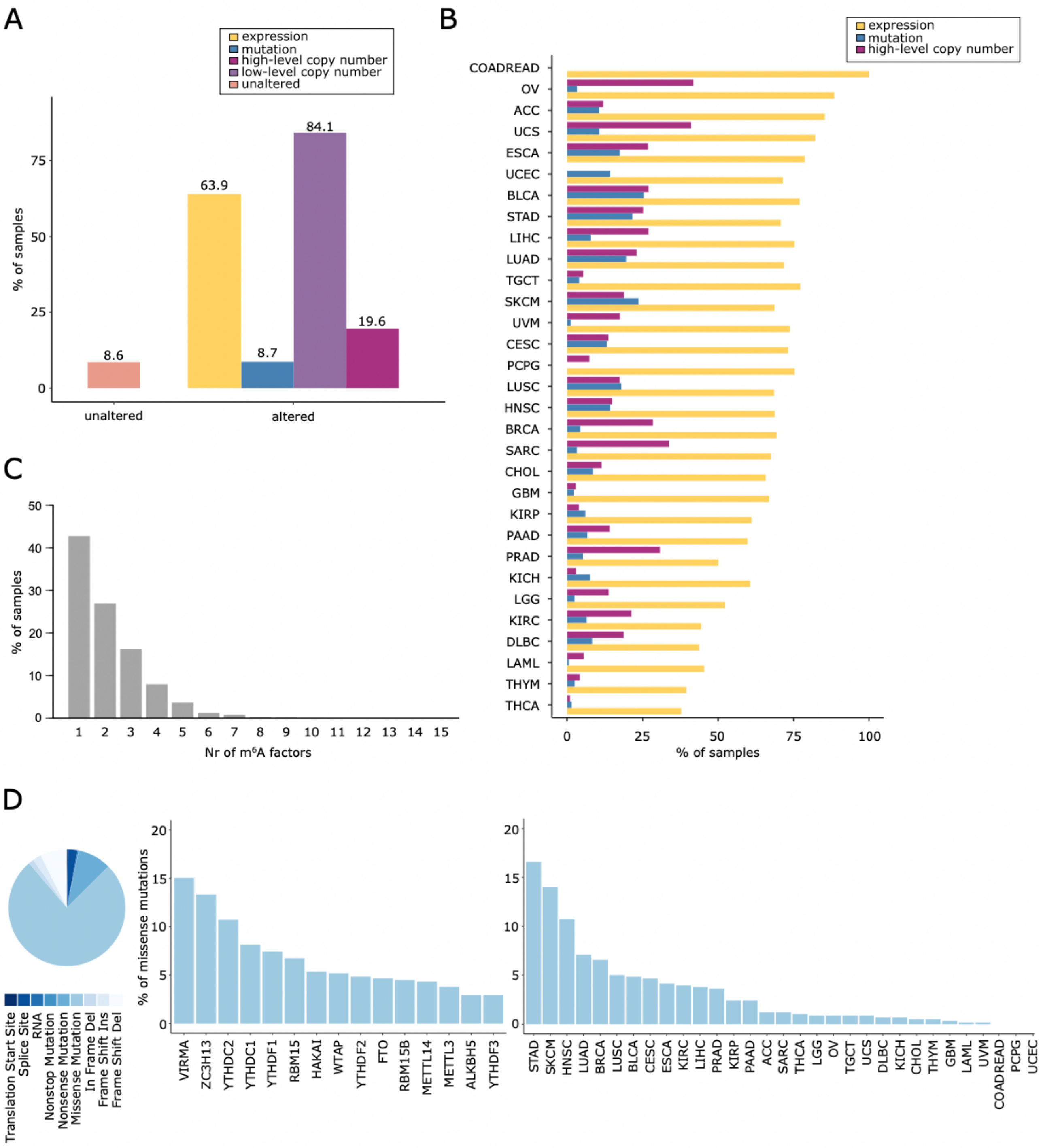


**Supplementary Figure 1. Dysregulation of m$^6$A factor expression is present in 31 tumor studies, the majority of which have alterations in only one of the 15 m$^6$A core factors.** A) The plot shows the percentage of samples altered in m$^6$A factors stratified by presence/absence of high/low-level copy number, mutation, and/or expression alterations. B) The plot shows the percentage of samples with m$^6$A factors, high-level copy number, mutation, and/or expression alterations stratified by tumor types. C) The plot shows the number of samples with high-level copy number, mutation and/or expression alterations in one or more m$^6$A factors. D) Pie chart shows the proportion of mutation types. The bar plots show the percentage of missense mutations across m$^6$A factors and tumor types.

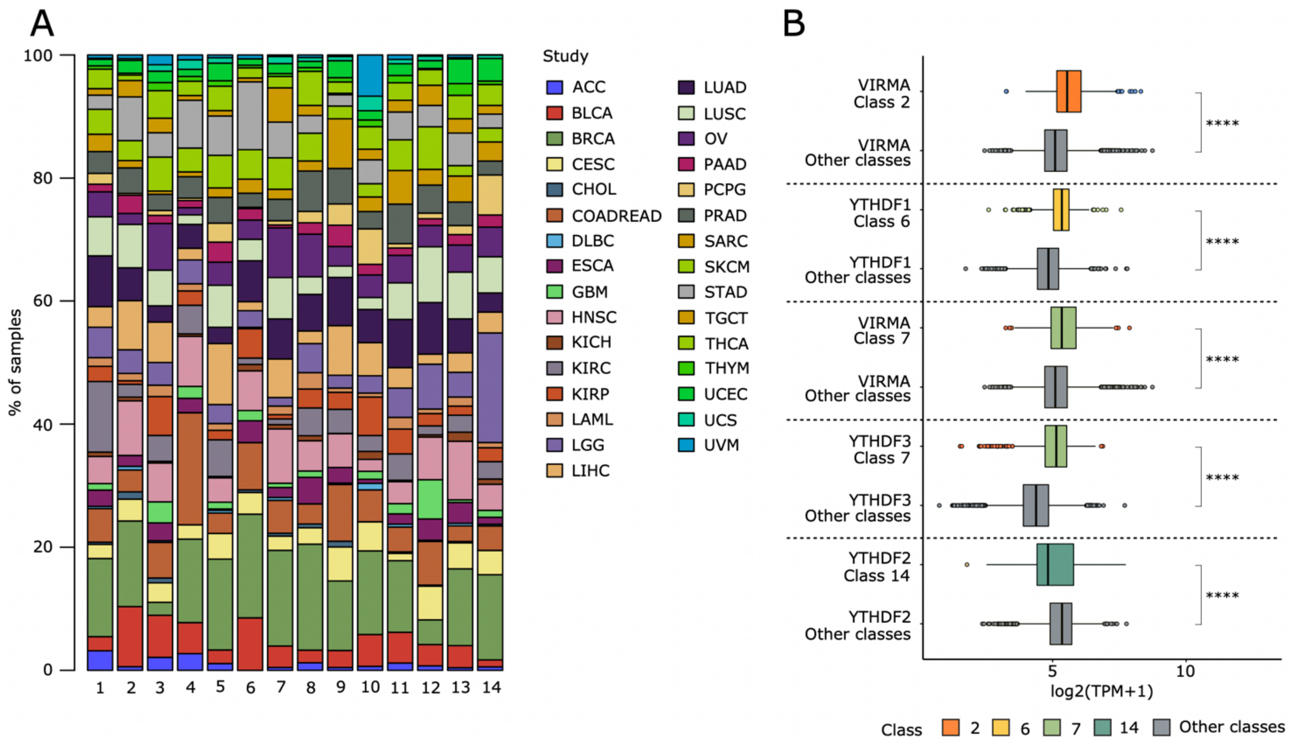


**Supplementary Figure 2. The most frequently dysregulated m$^6$A factors show altered expression levels in the classes with their unique alteration, which are enriched for 5 tumor types.** A) Tumor type composition of the 14 classes. B) The boxplots show the comparison between the expression (Xena portal - TPM expression) of the most frequently altered m$^6$A factors (VIRMA, YTHDF1, YTHDF2, and YTHDF3) in the four respective classes (classes 2, 6, 7, and 14) and the remaining classes. Wilcoxon test: ****p<=0.0001, ***p<=0.001, **p<=0.01, *p<=0.05.

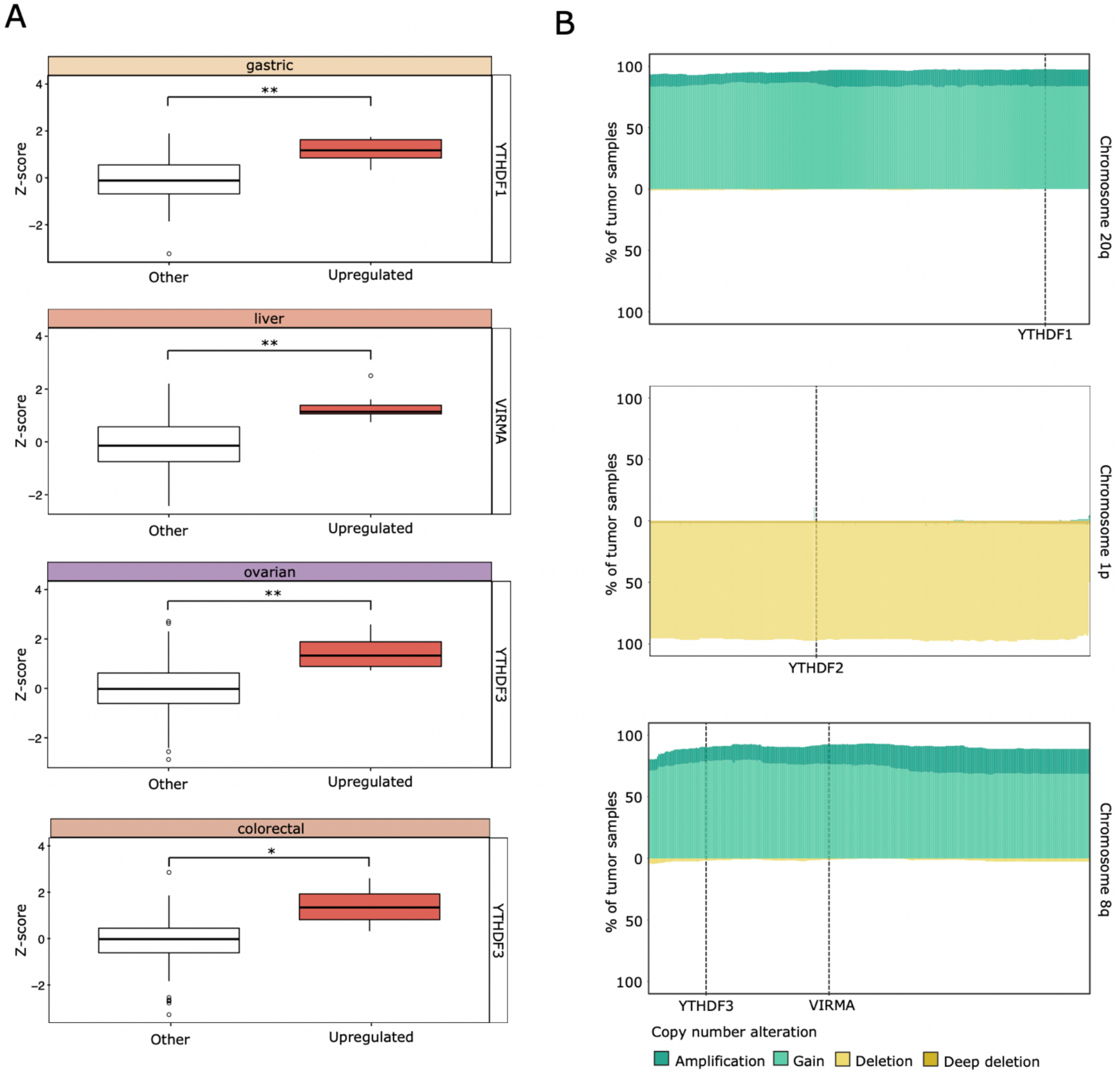


**Supplementary Figure 3. Tumor types with alterations in the most frequently dysregulated m$^6$A factors show associations between mRNA-level dysregulation and protein expression and frequent chromosomal arm copy number changes.** A) The protein levels of the most frequently altered m$^6$A factors are shown for the available tissues corresponding to the tumor types enriched in the dysregulated expression of the respective m$^6$A factors. In each plot, the samples were categorized based on the mRNA expression (upregulation: z-score >= 2), and the protein levels were compared between upregulated and the remaining samples. Wilcoxon test: ****p<=0.0001, ***p<=0.001, **p<=0.01, *p<=0.05. B) The plots display the percentage of samples with CNAs in the genes located on the chromosome arms of the most frequently altered m$^6$A factors (YTHDF1-chr20q, YTHDF3/VIRMA-chr8q, YTHDF2-chr1p). The names of the genes are hidden from the x axis apart from the considered m$^6$A factors. Each plot represents samples from tumor types enriched in the dysregulation of the expression of the corresponding m$^6$A factor.

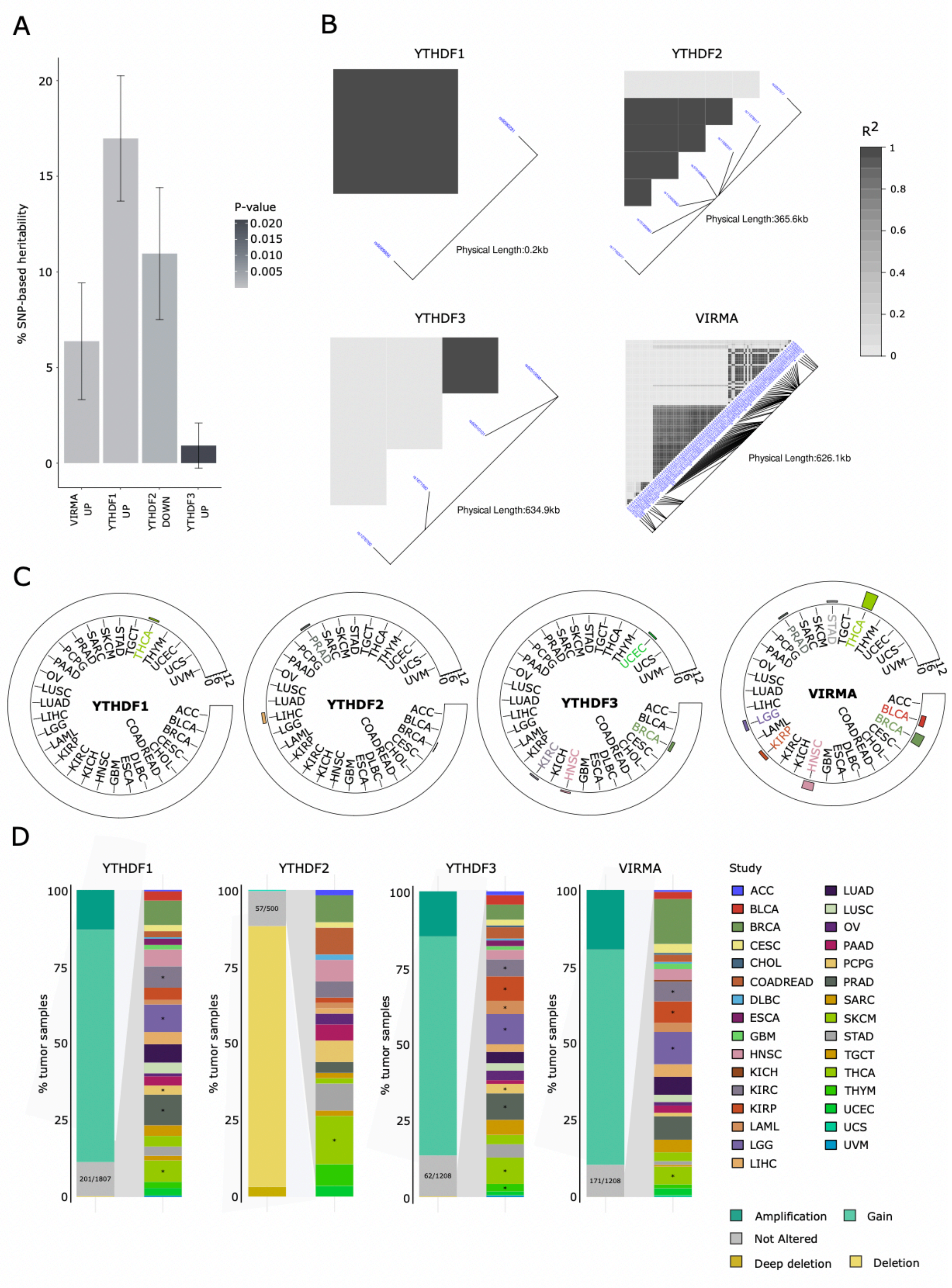


**Supplementary Figure 4. eQTLs may play a role in driving alterations in the most frequently dysregulated m<sup>6</sup>A factors, in the absence of copy number alterations.** A) The four cancer-associated phenotypes (VIRMA upregulation, YTHDF1 upregulation, YTHDF2 downregulation, and

YTHDF3 upregulation) show significant levels of SNP-based genome-wide heritability (V(Genotype)/Vp) (LRT p-value < 0.05). B) Linkage disequilibrium (LD) plots generated using the LDheatmap R package. The relative physical position of each SNP is given in the lower diagram, and the pairwise linkage disequilibrium (R2) between all SNPs is given above each SNP combination. C) The plots display the number of strong LD groups in each tumor type. D) The plots show the percentage of samples with expression dysregulations in the most frequently altered $m^6A$ factors. The number of samples with expression dysregulations and without collinear copy number alterations (shown in grey) are displayed along with the corresponding percentages of these samples within each tumor type. Enrichment analysis was performed with Fisher exact test: * adjusted p-value <= 0.05 and odds ratio >= 3.

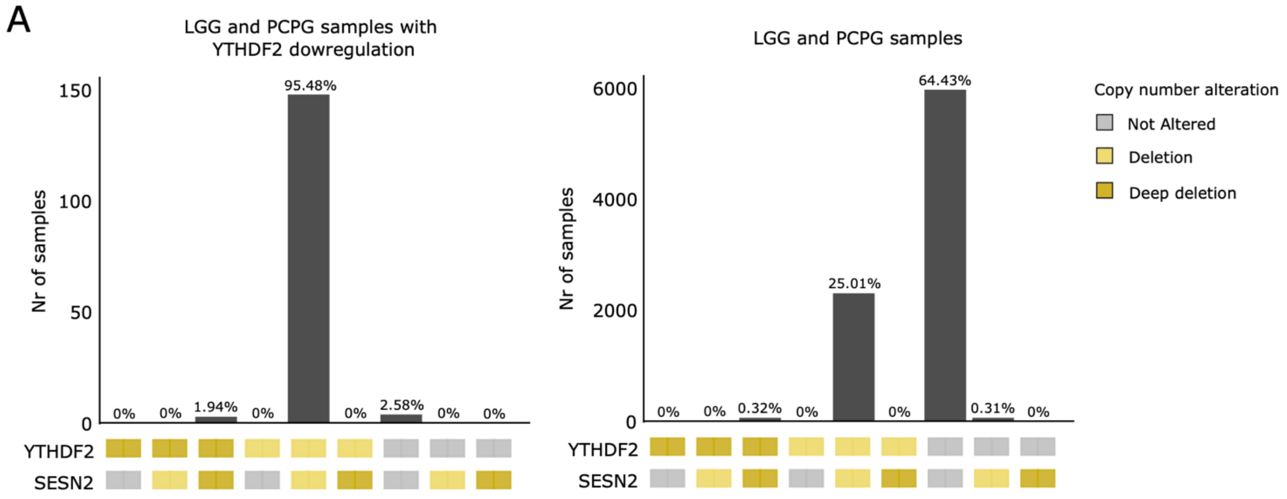


**Supplementary Figure 5. Hitchhiking effects of the m$^6$A factors by near cancer driver genes is excluded.** A) The two plots show the deletion patterns of YTHDF2 and SESN2 genes in the tumor types enriched in YTHDF2 downregulation (LGG, PCPG). On the left, only samples with YTHDF2 downregulation were considered, and on the right all the samples.

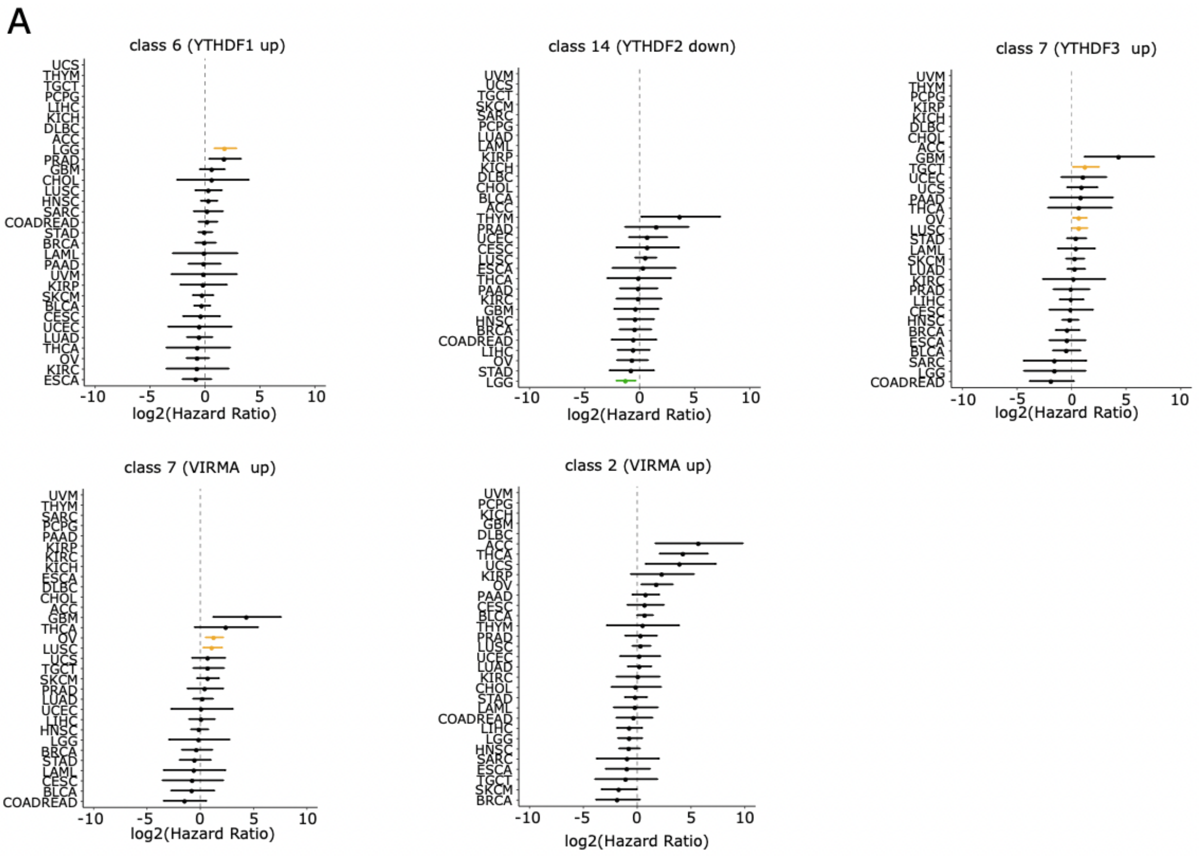


**Supplementary Figure 6. The most frequently dysregulated m$^6$A genes show single tumor prognostic roles in their respective classes.** A) The forest plots show the results of multivariate cox regression analyses performed on samples of single tumor types of class 6 with YTHDF1 upregulation, of class 14 with YTHDF2 downregulation, of class 7 with YTHDF3 upregulation, and of class 7 and 2 with VIRMA upregulation. Yellow/green tumor studies are significantly associated with a worse/better prognosis (adjusted p-value < 0.1).
